# Bootstrapping and Empirical Bayes Methods Improve Rhythm Detection in Sparsely Sampled Data

**DOI:** 10.1101/118521

**Authors:** Alan L. Hutchison, Ravi Allada, Aaron R. Dinner

## Abstract

**Motivation:** There is much interest in using genome-wide expression time series to identify circadian genes. However, the cost and effort of such measurements often limits data collection. Consequently, it is difficult to assess the experimental uncertainty in the measurements and, in turn, to detect periodic patterns with statistical confidence.

**Results:** We show that parametric bootstrapping and empirical Bayes methods for variance shrinkage can improve rhythm detection in genome-wide expression time series. We demonstrate these approaches by building on the empirical JTK_CYCLE method (eJTK) to formulate a method that we term BooteJTK. Our procedure rapidly and accurately detects cycling time series by combining information about measurement uncertainty with information about the rank order of the time series values. We exploit a publicly available genome-wide dataset with high time resolution to show that BooteJTK provides more consistent rhythm detection than existing methods at typical sampling frequencies. Then, we apply BooteJTK to genome-wide expression time series from multiple tissues and show that it reveals biologically sensible tissue relationships that eJTK misses.

**Availability:** Bootstrap eJTK (BooteJTK) is implemented in Python and is freely available on GitHub at https://github.com/alanlhutchison/BooteJTK.

## 1 Introduction

Periodic patterns (rhythms) are pervasive in biology at molecular, cellular, organismal, and ecological scales. It can be challenging to detect these patterns genome-wide with confidence of their significance, however, because biological dynamics are intrinsically noisy, and often it is feasible to obtain only a few samples of a potentially periodic process. To address this issue, several statistical methods have recently been developed to identify genes with cycling expression patterns from genome-wide time series [16, 30,4,36,14, 18].

Empirical JTK_CYCLE (eJTK) [14, 16] and RAIN [30] are non-parametric methods that analyze the rank order of measurements. While this approach makes them sensitive to waveforms of arbitrary shape, it does not incorporate information about the measurement uncertainty. ANOVA [18] is a parametric approach that does incorporate the variance of intra-time point measurements when identifying differences in mean values, but it is less sensitive than eJTK because it does not use information about the time order of the measurements [16]. ARSER [36] is likely the most successful parametric method at present, but the fact that it fits a time series to a sinusoidal curve by autoregressive spectral estimation makes it less sensitive to non-sinusoidal time series, as we discuss below.

Either explicit or implicit in these methods is comparison of the variation in measurements at each time point (i.e., across replicates/periods) to the variation from one time point to another over the period of interest. The cost and effort of sample preparation and measurement limits the number of replicates/periods obtained. As a result, the observed variation at each time point may poorly represent the true variance. In particular, if the data yield an estimate of the variation that is too small, a time series is more likely to be falsely identified as cycling because the apparent signal is large compared with the apparent noise. Properly accounting for small replicate numbers in estimating the variation has the potential to provide substantial gains in accuracy of rhythm detection and, in turn, aid in understanding periodic biological processes.

To this end, we introduce an empirical Bayes (eBayes) procedure [28, 21]. In this approach, which is commonly employed in differential expression analysis [31, 26], information from across a dataset is combined to estimate a prior distribution for the standard error, and this prior is then used together with the individual measurements to estimate the variance at each time point. This ‘shrinks’ the spread in variances (Fig. S1) [21]. Here, we use the empirical Bayes variance estimates to generate parametric bootstrap time series samples and then apply a rhythm detection algorithm to them. The parametric bootstrap [5] is also established in bioinformatics, and is applied in packages for RNA-Seq quantification [2] and differential expression [25]. To the best of our knowledge, this is its first application in rhythm detection.

While the strategy is general, we focus on its implementation with the empirical JTK_CYCLE (eJTK) method, which we have demonstrated to outperform most other algorithms [16].eJTK compares time series to a set of reference waveforms varying in phase (peak expression) and distance from peak to trough using a non-parametric rank order correlation, Kendall’s τ. Selecting the best waveform presents a multiple hypothesis testing problem, which eJTK solves by empirically calculating the null distribution of the selection procedure to assign p-values to resulting rhythmicity scores. This approach is accurate but relatively computationally costly because the null distribution must be re-evaluated for each set of measurements (in order to distinguish time series that are missing observations for different sets of time points). In the present work, we reduce this expense significantly by fitting a Gamma distribution to test statistics for a small number of time series. This approximation makes eJTK, even in the context of the bootstrap, computationally economical.

Our approach, which we term BooteJTK, combines freedom from restrictive assumptions regarding the shape of the waveform with incorporation of information about the uncertainty in each measurement. We demonstrate the method on simulated data and two circadian genome-wide expression datasets. The first dataset is densely sampled with measurements of gene expression in mouse liver samples at 1 h intervals for 48 hours [13]; this dataset allows us to examine the performance (self-consistency) of the method as fewer time points are included. The second dataset comprises gene expression measurements every 2 h for 48 hours for 12 mouse tissues in continuous darkness [37]. This dataset allows us to look at the consistency of rhythm detection across tissues.

We find that fewer genes are rhythmic than previously believed, due to the more stringent requirement that the uncertainty in measurements be small relative to the amplitude of expression, in addition to the rank order of the values of the time series matching those of a reference waveform. Corroborating our more stringent results with core clock transcription factor targets (CCT) [20], we find no decrease in CCT enrichment between BooteJTK and eJTK. At the same time, we find increases in the overlap of rhythmic genes across tissues, suggesting circadian programs in different tissues may not be as distinct as previously thought. Put together, the results indicate that BooteJTK provides robust rhythm detection with improved consistency. The general principles and methods that we present here, the empirical Bayes and bootstrapping procedures, can be applied to other rhythm detection algorithms.

## 2 Methods

Our method, BooteJTK, first obtains time point mean and standard deviation estimates by empirical Bayes variance estimation from a time series. It then runs eJTK on parametric bootstrap replicates of the time series, averaging the rhythmicity scores to obtain a new rhythmicity score, which is assigned a p-value. These steps can be seen as a flow chart in Figure S2.

### 2.1 Empirical Bayes variance estimation

Empirical Bayes methods are an established part of many workflows for differential expression analysis [28, 31, 26]. These methods combine information from all the time points in the data to ‘shrink’ the spread of standard deviation estimates. As a result, low standard deviation estimates are increased and high standard deviation estimates are decreased (Fig. S1). Given that we have low replicate numbers, it is beneficial to use empirical Bayes methods. In particular, we use *voom* (specifically, *vooma* for microarrays) [26] to obtain initial estimates of the standard deviation that take mean-variance relationships into account. These estimates are then adjusted using the emprical Bayes procedure implemented in *vash* [21]. We originally used *limma* [26] in place of *vash,* but that resulted in over-dispersion and over-estimates of rhythmicity p-values, partially due to adjustment of small standard errors away from zero.

### 2.2 Bootstrapping eJTK

In bootstrapping, data are resampled with replacement to create a distribution of simulated measurements that can in turn be used to compute statistics [5]. Genome-wide circadian time series generally consist of two or three replicates per time point, minimizing the benefits offered by resampling the data directly (i.e., non-parametric bootstrapping). Instead, we use parametric bootstrapping. Specifically, we log-transform the expression measurements [32] and model the resulting data at each time point as normally distributed with the mean directly calculated from the replicates and the variance modeled by *voom* [26] and the empirical Bayes procedure *vash* [21], both implemented in R (Fig. S1).

We generated time series for this model and analyzed them with eJTK to determine their circadian characteristics: rhythmicity score (τ), phase (peak), and best-matching waveform. We averaged each of these statistics across the model time series. While eJTK generally outputs integer multiples of the measurement interval for the peak and trough times (i.e., extrema), the means of these statistics can be non-integer, which allows for better representation of the times of the extrema when they do not coincide with the measurement times. Regardless, for the phase and trough, the mean values are close to the values output by eJTK. This is not necessarily the case for the rhythmicity score, as we now discuss.

In the context of eJTK, the Kendall’s τ statistic measures the correlation in rank order of the values of the time series of interest and the values of a discretized reference waveform; the rhythmicity score is the highest τ across all tested reference series. A perfect match in rank order has τ = 1. Adding noise to the values of a reference time series and comparing the resulting rank order with the original one often results in τ < 1, with τ tending to decrease as the noise becomes larger in comparison with the amplitude of the oscillation. As a result, the mean of the distribution of τ values for the bootstrap resamples depends on both the rank order of time series values and the measurement uncertainty.

An additional issue is that the τ distribution is skewed when τ is close to the limits of its range (-1 and 1). To stabilize the variance across the full range of possible rhythmicity scores, we average the Fisher transform of τ: τ̃=arctanh(τ), truncating the values to ±arctanh(0.99) for ±τ > 0.99 to ensure that the τ̃ values are finite.

### 2.3 Obtaining Accurate and Computationally Inexpensive P-values

A p-value is the likelihood under the null hypothesis of observing a value of a test statistic or a more extreme one. We previously generated the null distribution for eJTK by applying the method to 10^6^ time series generated by selecting values from a Gaussian distribution with a constant mean [16]. We repeated this procedure for each number of measured time points in the experimental time series. Because this numerical procedure represents most of the computational expense of eJTK and, in turn, BooteJTK, we sought an approximate analytical form for the null distribution. We found the Gamma distribution, which we previously used to model the F24 null distribution [16], to be a reasonable choice (Figs. 1 and figS3). To assess this approach quantitatively, we computed p-values with this model and our earlier method (i.e., empirically from the histogram of 10^6^τ̃values) for a range of τ̃ values. The ratio of the Gamma-distribution-generated p-values to the empirical p-values is larger than 1 at very low p-values and close to 1 for moderate p-values close to typical significance thresholds (figs. 1B and S3B). This means that the p-values obtained from the Gamma-distribution approximation are sufficiently accurate for use, but they favor the null hypothesis relative to the empirical values, leading to slightly fewer genes being considered rhythmic for a given significance threshold.

**Figure 1:**
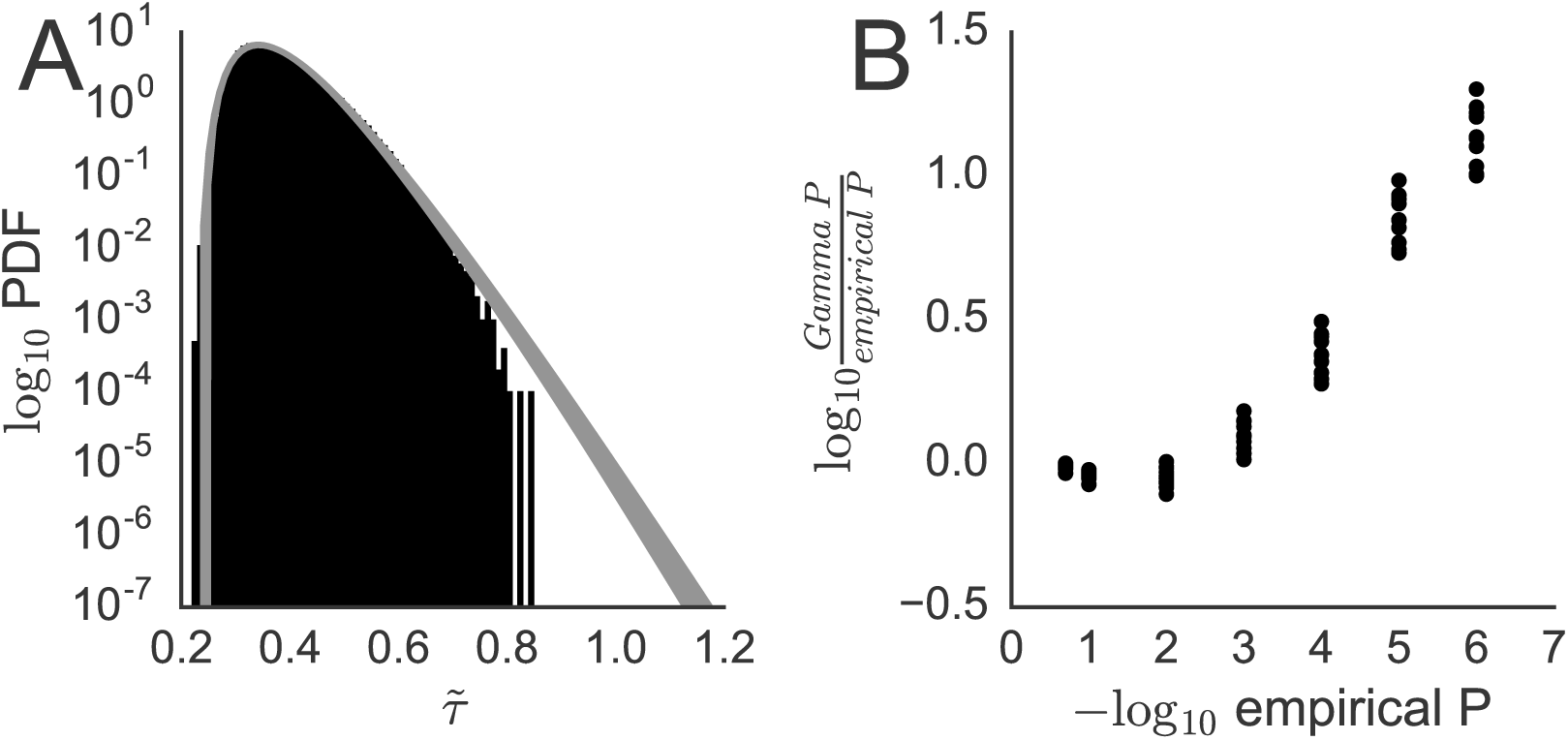
The BooteJTK τ̃ null distribution can be modeled by a Gamma distribution. (A) Comparison between Gamma distributions parameterized by the Python scipy.stats.gamma.fit [17] function of τ̃ values for 10^3^ time series comprised of 24 time points drawn randomly from a Gaussian distribution and a histogram of 10^6^ τ̃values from similarly generated time series. 10 different such Gamma distributions are shown. (B) Logarithms of the ratios of the p-values estimated from the Gamma distributions in (A) (Gamma P) to the p-value calculated directly from the cumulative distribution function for the 10^6^ τ̃ values used to construct the histogram (empirical P).

### 2.4 BooteJTK Outperforms Alternative Rhythm Detection Methods

To test the ability of BooteJTK to identify rhythmic time series, we generated 100 time series from a cosine with a 24 h period sampled every 2 h in duplicate with Gaussian noise added to each point. We varied the Gaussian standard deviation relative to the amplitude of the underlying cosine, a ratio we refer to as the noise level. To model the null hypothesis, we generated 1000 time series with the same number of time points but with values drawn only from the Gaussian distribution.

#### 2.4.1 BooteJTK outperforms eJTK

We compared BooteJTK to eJTK, approximating the null distribution as a Gamma distribution in both cases. In each case, we compared the simulated time series against cosine reference waveforms with 24 h periods, testing phases every 2 h from 0 h to 22 h and asymmetries every 2 h from 2 h to 22 h (132 total reference waveforms), as described in Hutchison *et al.* [16]. Figure 2 shows that BooteJTK outperforms eJTK: for different noise levels the True Positive Rate (TPR) is higher fora given False Positive Rate (FPR) (Figs. 2A and S4A), and the Matthews Correlation Coefficient (MCC) is higher for all p-values (Fig. 2B and S4B). The MCC is 1 if a classifier is perfect and 0 if it performs no better than random guessing.

**Figure 2:**
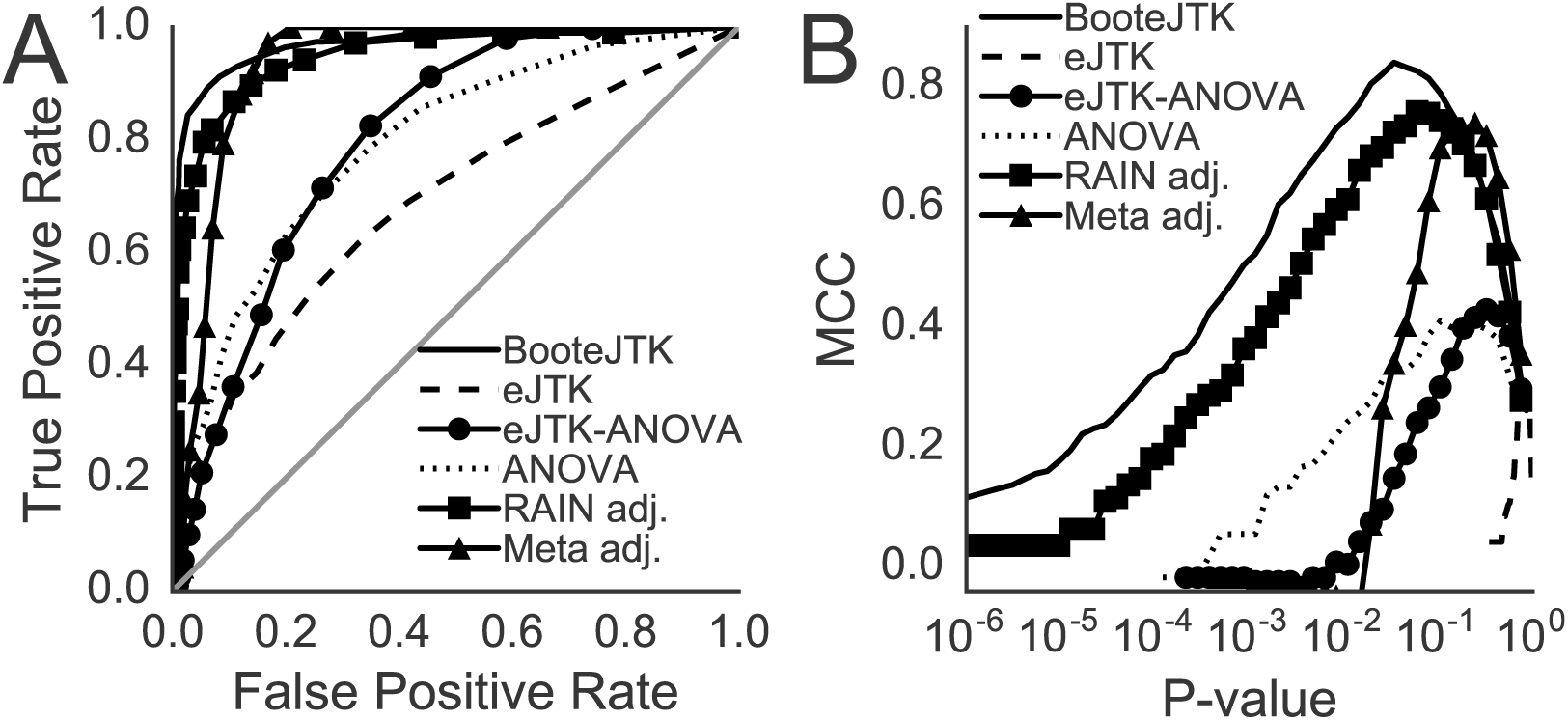
BooteJTK outperforms alternative methods in detecting simulated sinusoidal time series as measured by (A) True Positive Rate (TPR) against False Positive Rate (FPR) and (B) Matthews Correlation Coefficient. As indicated in the legend, we compare BooteJTK with 25 bootstrap samples to adjusted RAIN [30, 15], adjusted MetaCycle [34, 15], empirical JTK (eJTK) [16], ANOVA [18], and a Brown-Fisher method p-value integration [6, 3] of eJTK and ANOVA (eJTK-ANOVA). Simulated data are generated with noise-to-amplitude ratio of 1.00.

To gain a better sense of the differences between BooteJTK and eJTK, we selected two time series from the simulated data with noise level 0.60 that had the same τvalue for eJTK(τ̃= 0.57, p = 0.002) but different values for BooteJTK (τ̃= 0.67 and 1.08, *p* = 0.008 and 0.001) (Figs. S5A and B). Using the Benjamini-Hochberg correction to control for the false discovery rate when testing many time series [1], the first time series would likely be considered arrhythmic, while the second time series would likely be considered rhythmic. In Fig. S5A, the average time series matches the original well, but the standard deviations around the mean points are large and overlap substantially. The overlap results in a lower BooteJTK score. In Fig. S5B, the original time series matches the average and the standard deviations around the mean points are small and distinct from one another. The tighter uncertainties result in a higher BooteJTK score.

#### 2.4.2 BooteJTK outperforms RAIN

We also compared BooteJTK to RAIN [30], another non-parametric method that uses reference waveforms. RAIN does not take into account the size of the noise relative to the amplitude of the time series, which we expect to be more important when testing experimental data. We had previously found that RAIN underestimates p-values, so we adjusted RAIN using 10^6^ simulations of the null distribution as described in Hutchison *et al.* 2017 [15]. We found that BooteJTK outperforms RAIN as measured by TPR vs. FPR and by MCC curves (Fig. 2).

#### 2.4.3 BooteJTK outperforms ARSER and ANOVA on asymmetric time series

As mentioned in the Introduction, parametric methods do account for the size of the noise relative to the amplitude of the time series, and ARSER [36] is likely the best such method presently. Because ARSER fits a time series to a sinusoidal curve, we expect it to outperform nonparametric methods when detecting time series that are approximated well by that waveform. However, we expect many biological time series to be non-sinusoidal [16]. For this reason, we compared BooteJTK, ARSER, and a reference free parametric method, ANOVA [18], for their abilities to detect 24 h time series with peak to trough intervals of 20 h before adding noise. BooteJTK outperforms both methods as shown by TPR vs. FPR and MCC curves (Fig. S4C and D).

#### 2.4.4 BooteJTK outperforms MetaCycle and combined eJTK-ANOVA

We observed that BooteJTK combines eJTK’s non-parametric aspects with ANOVA-like comparison of the relative measurement uncertainty (variance between replicate points) to the amplitude of the entire time series (variance across the time series). We thus wondered if a combination of ANOVA and eJTK p-values would perform similarly to BooteJTK. A new method, MetaCycle, was recently developed that combines ARSER, the original JTK_CYCLE, and Lomb-Scargle by integrating their p-values using Fisher integration to increase rhythm detection ability [6, 34]. We showed that their method underestimates p-values and can be improved by using the Brown correction of the Fisher integration method [3] as well as empirically calculating their p-values [15]. We used Fisher integration with the Brown correction to combine ANOVA and eJTK and compared it to BooteJTK. BooteJTK outperformed both eJTK-ANOVA and adjusted MetaCycle (Fig. 2). The combination of eJTK and ANOVA may not be suitable for improved rhythm detection and this result affirms that the classification strength of BooteJTK is greater than a combination of eJTK and ANOVA.

### 2.5 Computational Expense

Having established the sensitivity and specificity of BooteJTK, we sought to minimize the computational cost of our method. For the simulated data described above, we found no discernible difference in τ̃scores for 100, 50, 25, and 10 bootstrap samples (Fig. S6). We use 25 bootstrap samples throughout the rest of this study. With 25 bootstrap samples, we obtain run times on a Late-2013 iMac Desktop with a 3.5 GHz Intel Core i7-4771 processor and 16 GB of 1600 MHz DDR3 memory. For 1000 time series, the analysis by BooteJTK took 180 s and the analysis by eJTK took 8 s. Of this, less than 2 s is the integration of the Gamma distribution to translate τ̃statistics to p-values. With the improvements in the present paper, the computational cost of eJTK is several orders of magnitude less than in Hutchison *et al.* 2015 [16].

## 3 Results

### 3.1 Effect of sampling frequency on rhythm detection

While BooteJTK outperforms leading methods for simulated data, such time series can lack features of experimental data. Assessing the behavior of algorithms for experimental data can be challenging, however, because rhythmic expression has been independently verified for only a small fraction of the genome. Nevertheless, we expect rhythm detection to be accurate when a waveform is extensively sampled (with high frequency, over many periods), and we can study the consistency of each method as data are downsampled.

To this end, we applied BooteJTK to microarray data collected every 1 h for 48 h from mouse liver tissue under constant conditions [13]. As the original analysis of the dataset was performed with JTK_CYCLE [14], we analyzed the dataset with eJTK as well for comparison. We treated the modulo 24 time points as replicates, providing 2 replicates every 1 h over 24 h. Since most transcriptomic circadian experiments have data collected every 2 h [37] or 4 h [8, 24], we parsed the data (from CT18-CT65) into two datasets with measurements every two hours (denoted 2a: CT18, CT20, etc. and 2b: CT19, CT21, etc.) and into four datasets with measurements every four hours (denoted 4a, 4b, 4c, and 4d, starting at CT18, CT19, CT20, and CT21, respectively).

Using the R package *gcrma* [35] to normalize the data (GEO GSE11923) and removing probes with constant expression, we compared the Benjamini-Hochberg adjusted p-values (BH) from BooteJTK to eJTK on the full dataset using a significance threshold of 0.05 (Fig. 3A). BooteJTK identifies more probes as rhythmic than eJTK does. When BooteJTK identifies a probe as rhythmic, it tends to assign it a smaller p-value (i.e., greater significance) than eJTK, which leads to smaller Benjamini-Hochberg adjusted p-values. BooteJTK can provide smaller p-values because it checks whether the values of a time series have the right rank order (the sole criterion for eJTK) *and* whether the differences between pairs of points are large compared to the uncertainties in those measurements. The additional requirement makes it harder for a rhythmic pattern to arise by chance.

**Figure 3:**
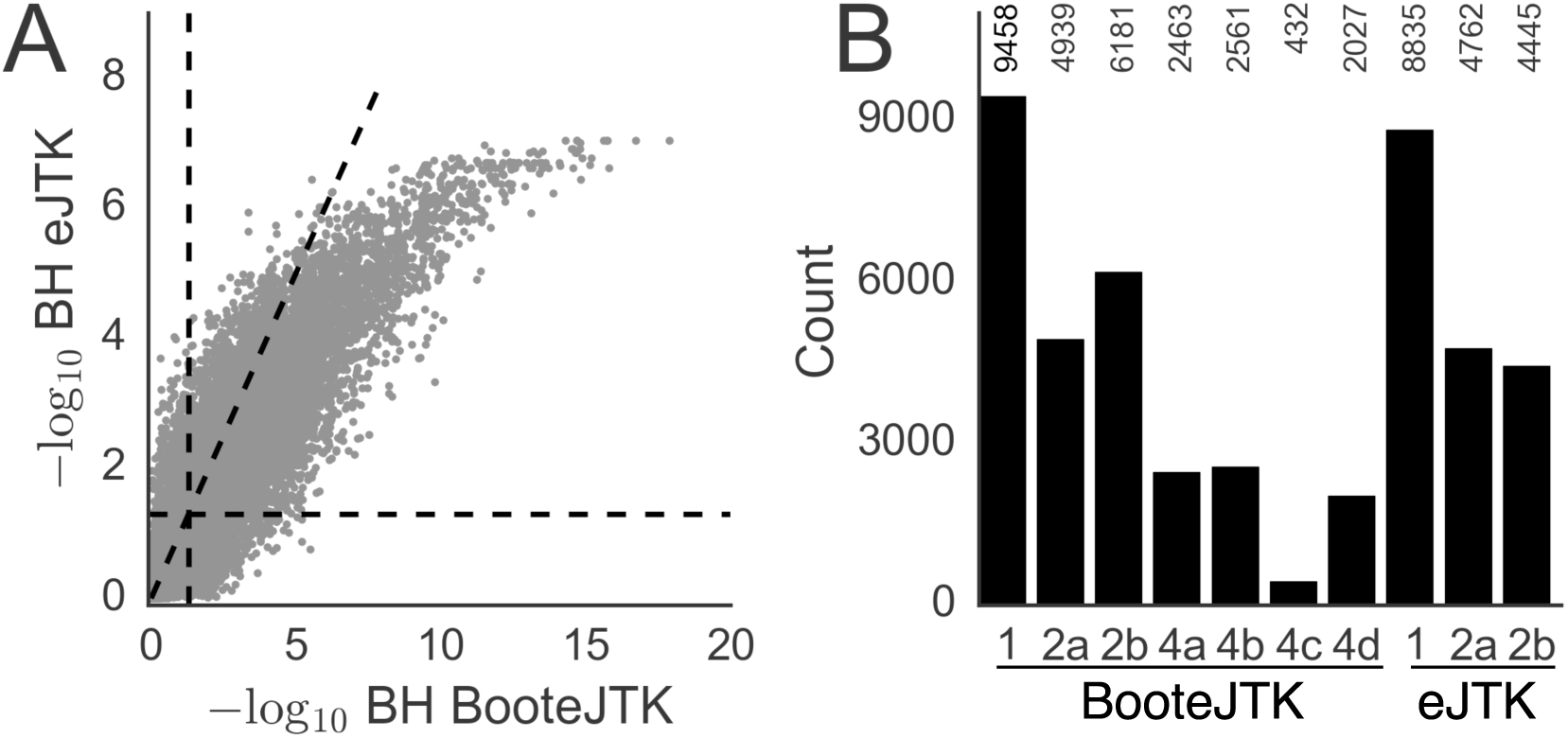
BooteJTK identifies rhythmic probes more consistently than eJTK as data are downsampled. Data shown are from Hughes *et al.* [13] and are originally sampled every 1 h for 48 h; the datasets are downsampled to measurement intervals of 2 h (denoted 2a and 2b) and 4h (denoted 4a, 4b, 4c, and 4d). (A) Comparison of BooteJTK and eJTK BH values for the full dataset. The diagonal line has a slope of 1; horizontal and vertical lines indicate–log_10_(BH = 0.05) ≈ 1.3. (B) Number of rhythmic probes at BH < 0.05 for the indicated methods and datasets downsampled to measurements every two hours (denoted 2a: CT18, CT20, etc. and 2b: CT19, CT21, etc.) and to measurements every four hours (denoted 4a, 4b, 4c, and 4d, starting at CT18, CT19, CT20, and CT21, respectively); the full dataset sampled every hour is denoted 1. eJTK identifies zero probes as rhythmic when the data are downsampled by 4 h.

Downsampling the data to measurements every 2 h reduces the number of rhythmic probes identified by both methods using a BH-corrected p-value threshold of 0.05 (Fig. 3B), though a greater fraction of BooteJTK-identified probes remain. Downsampling the data to measurements every 4 h prevents eJTK (and adjusted RAIN, Fig. S7) from finding any rhythmic probes at this significance threshold, while BooteJTK identifies thousands of the probes that it originally considered rhythmic. As noted above, BooteJTK incorporates more information, and the results are thus less sensitive to Benjamini-Hochberg correction.

The overlap between results obtained from two downsampled datasets can be quantified by the probability that a probe is rhythmic in one dataset (a row in Fig. 4B) if it is rhythmic in another (a column in Fig. 4B). For example, the probabilities are 0.78 and 0.62 ((row 3, column 2) and (row 2, column 3), respectively) that BooteJTK considers a time series downsampled to measurements every 2 h to be rhythmic if it identified that time series as rhythmic in the other dataset downsampled to measurements every 2 h. For eJTK, these values are 0.63 and 0.68 ((9,10), and (10,9), respectively) and for RAIN they are 0.61 and 0.68 (Fig. S8, (12,13) and (13,12), respectively). These results indicate greater consistency for the BooteJTK results and provide insight into how much inconsistency we should expect from experimental uncertainty because no biological differences should exist between downsampled sets.

**Figure 4:**
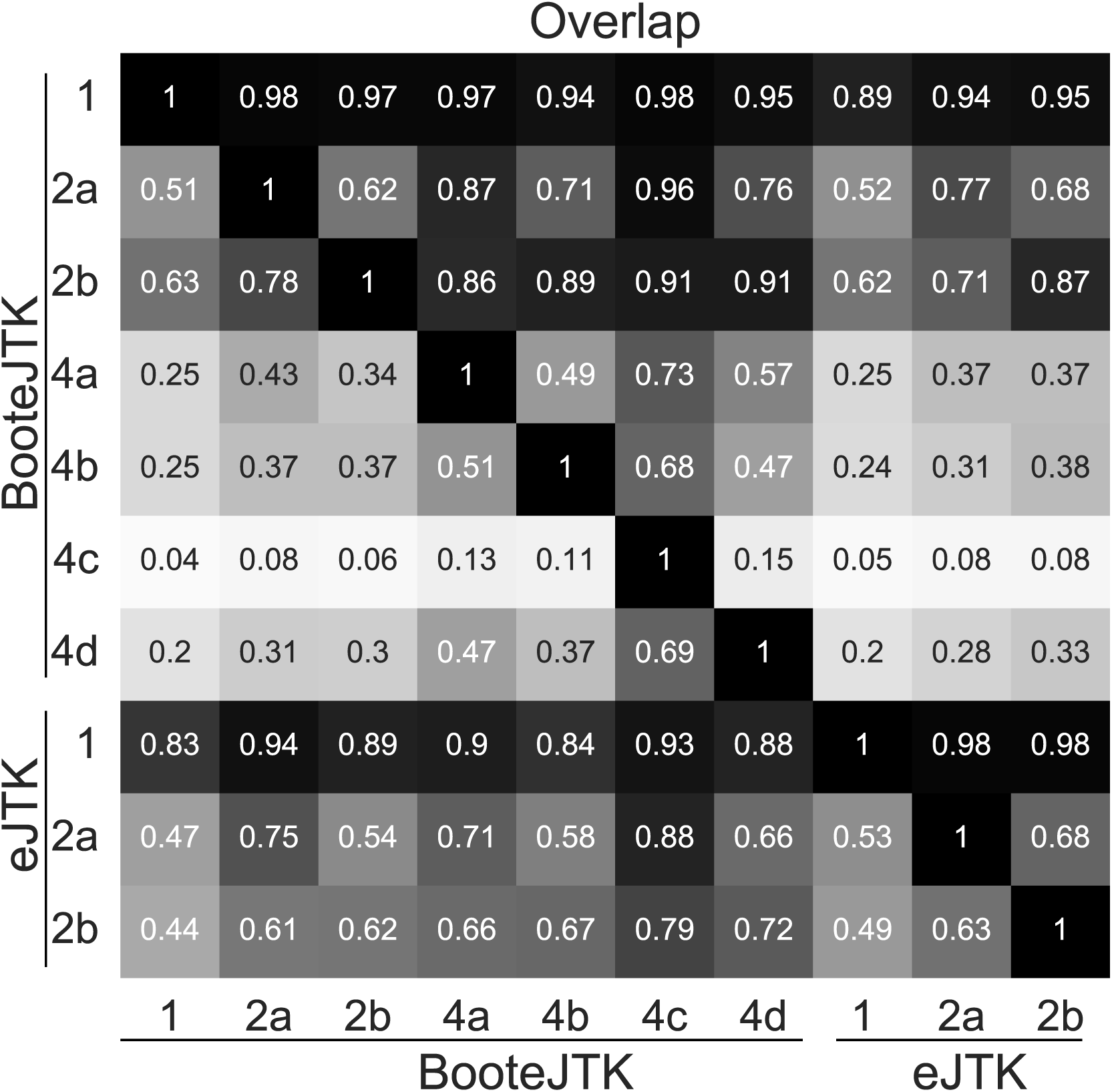
BooteJTK provides more consistent rhythm detection than eJTK between downsampled datasets. We quantified the overlap between results with different levels of downsampling by the probability that a probe is rhythmic in one dataset (a row) if it is rhythmic in another (a column). As no probes are found to be rhythmic when using eJTK on data downsampled to every 4 h, rows and columns for those datasets are not shown.

### 3.2 BooteJTK Reveals More Biologically Consistent Circadian Rhythmic Gene Expression Across 12 Mouse Tissues

Having established the improved rhythm detection of BooteJTK, we analyzed the 12-tissue mouse microarray time series in Zhang *et al.* [37]. The twelve tissues are adrenal (Adr), aorta, brown fat (BFAT), brainstem (BS), cerebellum (Cere), heart, hypothalamus (Hypo), kidney, liver, lung, muscle (Mus), and white fat (WFAT). Expression was sampled every 2 h for 48 h, which we again treated as duplicate measurements over 24 h. We also applied eJTK, mirroring the application of the original JTK_CYCLE method [14] by Zhang *et al.* [37].

### 3.2.1 BooteJTK is more stringent than eJTK but exhibits comparable enrichment for core clock targets

When applied to the 12-tissue microarray dataset from Zhang *et al.* [37], BooteJTK identifies 14,598 out of 25,268 probes (11,731/20,038 genes) as rhythmic in at least one tissue with BH <0.05, whereas eJTK finds 14,763 such probes (12,426 genes) (Figs. 5A and B). 12,261 of these BooteJTK-rhythmic probes are contained within the four tissues with the highest number of rhythmic genes: liver, kidney, lung, and brown fat (Fig. 5B). This difference was to be expected forthe reasons discussed above: BooteJTK compares the noise to the amplitude of a time series in addition to evaluating the rank order of the values. However, we were concerned that the greater stringency might exclude actual rhythmic time series. To evaluate this possibility, we corroborated our rhythmic genes with core clock transcription factor targets (CCTs) identified by ChIP-Seq in mouse liver [20]. Across the 12 tissues and between the two methods, we found no meaningful difference in the fraction of CCT genes relative to the number of genes identified as rhythmic (mean difference ‒0.016, standard deviation 0.022). This result persisted as we increased the number of TFs necessary to target a gene to qualify it as a CCT: the mean difference and standard deviation only became smaller with these added requirements.

**Figure 5:**
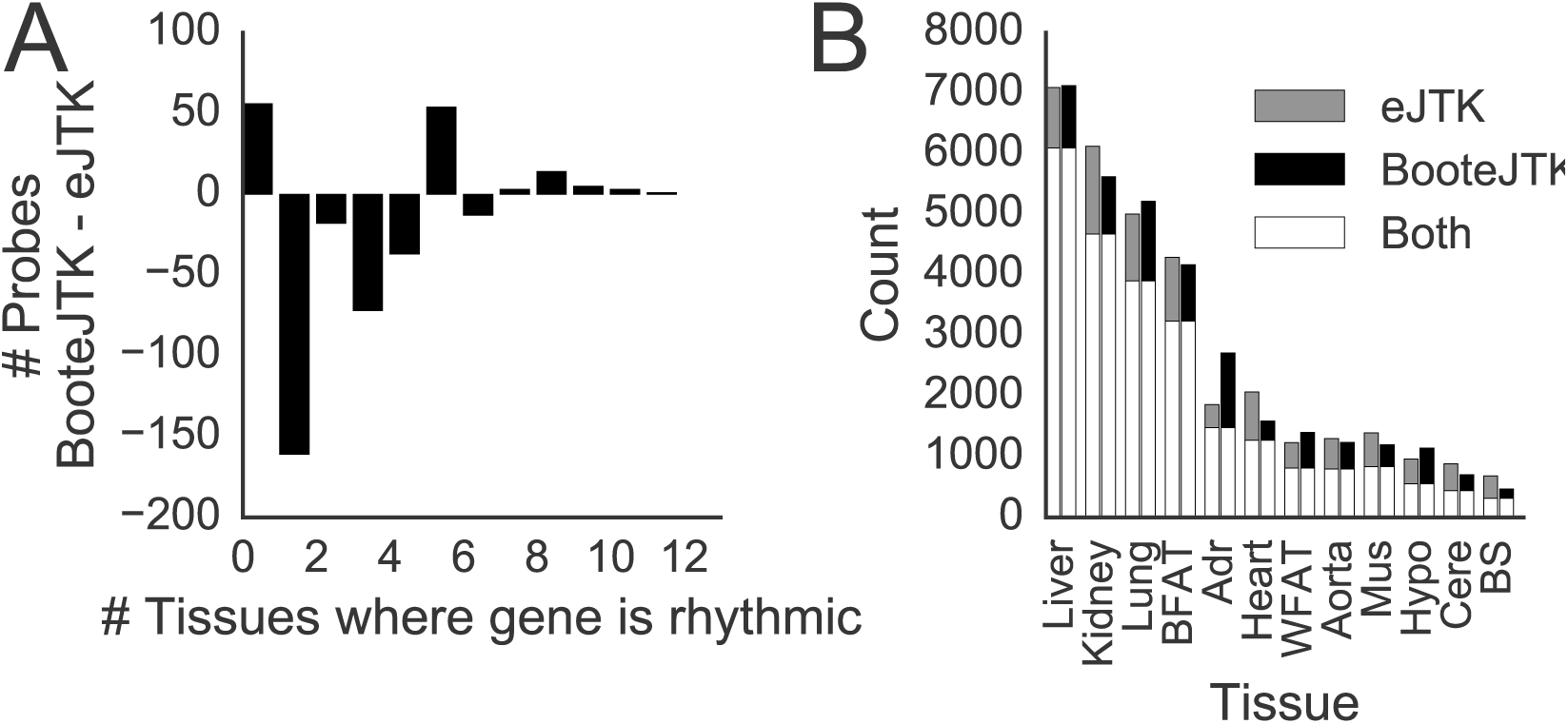
BooteJTK reveals fewer probes with circadian rhythmic expression across 12 mouse tissues than eJTK for the Zhang *etal.* dataset [37]. (A) Fewer probes are rhythmic in multiple tissues under BooteJTK than under eJTK (with BH < 0.05 for both methods). (B) Fewer probes per tissue are identified as rhythmic with BooteJTK than with eJTK.

### 3.2.2 Relationships between tissues

We examined the relationships between different tissues through the probability that a probe is rhythmic in one tissue conditioned on it being rhythmic in another (π). In Fig. 6A, row tissues are conditioned on column tissues, and the tissues are ordered by hierarchically clustering on the columns. Anatomically related tissues appear together in the plot—for example, the hypothalamus, brainstem, and cerebellum are on the right. It is important to note that the relationships between tissues are asymmetric: being rhythmic in the hypothalamus leads to a probability of 0.46 that a probe is rhythmic in the liver, but being rhythmic in the liver leads to only a probability of 0.07 that a probe is rhythmic in the hypothalamus. To put these numbers in context, we can compare them to the data of Hughes *et al.* [13]: the π values for two datasets from identical conditions downsampled to measurements every 2 h were 0.62 and 0.78. We note that a similarly high value is observed for the probability that a probe is rhythmic in brown fat conditioned on being rhythmic in the aorta (0.70). This result is interesting, but we feel further study is warranted as this point is an outlier in Fig. 6B, and the presence of brown fat around the aorta can easily lead to contamination of aorta samples [7].

**Figure 6:**
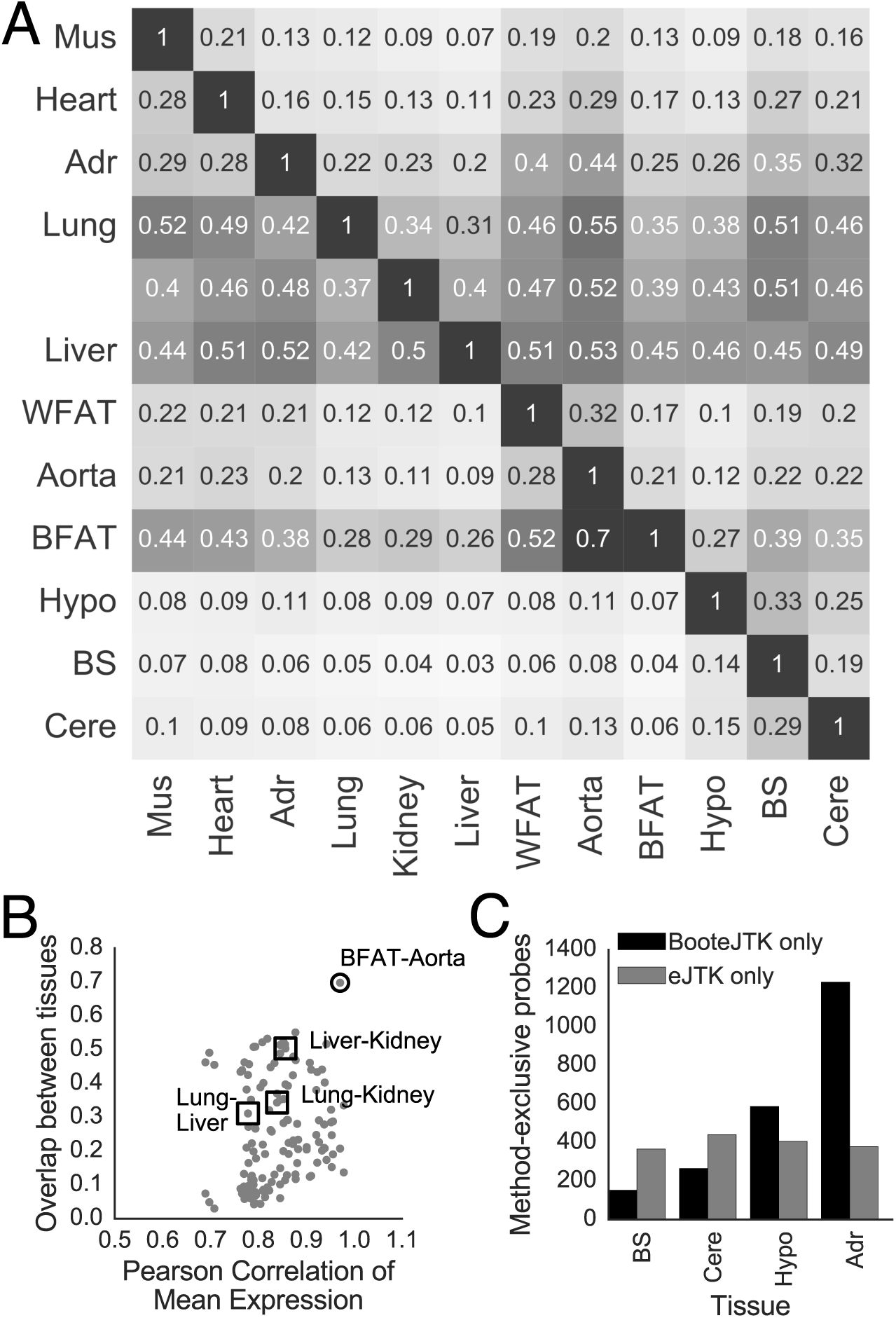
Overlap between tissues in rhythmic genes. (A) The probability that a probe is rhythmic in a row tissue if it is rhythmic in a column tissue (π) for BooteJTK. (B) Comparing correlation of mean probe expression between tissues with rhythmic overlap reveals that brown fat and aorta tissue have high correlation of gene expression and high rhythmic overlap relative to other tissue pairs, such as liver, kidney, and lung. (C) Numbers of rhythmic genes specific to each method for the tissues showing large differences in Fig. S9.

In Fig. S9, we show the differences between the π relationships from BooteJTK and eJTK. The columns corresponding to the brain tissues show marked differences. Zhang *et al.* [37] discuss the technical difficulty of dissecting the brain regions separately, so using a robust method to analyze these data should be of particular importance. The consistent increase in rhythmic predictive value of other tissues for the hypothalamus and adrenals is due to the increase in probes identified as rhythmic by BooteJTK relative to eJTK, whereas the increase in rhythmic predictive value ofthe brain stem and cerebellum regarding other tissues is due to a decrease in probes identified as rhythmic by BooteJTK relative to eJTK (Figs. 6C and S9).

### 3.2.3 Genes that are rhythmic in most tissues

Thirteen genes are identified as rhythmic by BooteJTK in all 12 tissues: Arntl (Bmal), Nr1d1 (Rev-erbA), Nr1d2 (Rev-erbB), Dbp, Per1, Per2, Per3, Ciart (Chrono), Bhlhe41 (Dec2), Tns2, Tsc22d3, Usp2, and Tspan4. Many of these are genes involved in the core clock machinery [33, 10]. However, known core clock genes such as Npas2, Tef, and Hlf are identified as rhythmic in only 11 of the 12 tissues; they do not meet the significance threshold in the hypothalamus. Given our prior knowledge regarding these genes and the evidence of their rhythmicity in other tissues, it is possible that they are in fact rhythmic in all 12 tissues and experimental issues are responsible for the inability to detect them in all 12 tissues. As noted above, Zhang *et al.* [37] suggest that the technical difficulty of dissecting the brain regions separately may negatively affect circadian rhythm identification in these tissues. We thus examined the 119 genes that BooteJTK identified as rhythmic in 9 or more of the tissues. Most of these genes were not rhythmic in the brainstem,hypothalamus, or cerebellum (Fig. S10).

Examining the functional annotation of these genes revealed many ontologies to be expected of consistently rhythmic genes, such as rhythmic processes and transcription regulation (Tab. 3.2.3). However, additional functional annotations were identified, such as genes involved in the stress response, endoplasmic reticulum, pigment granules, and heat shock. A few other genes stand out as well. Wee1 is rhythmic in 10 tissues (absent from hypothalamus and brain stem). Wee1 regulates cellular division by inhibiting entry into mitosis [19] and is known to be regulated by the core clock [22]; more generally, it has been suggested that the cell cycle is under circadian control [27]. Two other genes involved in the cell cycle, Cdkn1a and Calr, are rhythmic in 9 or more tissues. Given that many of these tissues have little cell proliferation, these genes may be functioning in other processes, in which case we expect those processes to be influenced by the circadian clock as well. Fmo1 and Gstt2 are identified as rhythmic in 10 tissues, while Fmo2 is identified as rhythmic in 11 tissues. These genes are identified as genes involved in drug metabolism by the DAVID webtool [11, 12]. Given the increasing interest in chronotherapeutics [37], further research into these genes is warranted in order to better understand their involvement in circadian processes.

## 4 Discussion

We have shown that rhythm detection from genome-wide expression time series can be considerably improved by using an empirical Bayes approach to improve variance estimates from limited replicates and propagating the resulting estimates into the test statistic for eJTK [16] by a parametric bootstrap. Because eJTK itself is nonparametric, BooteJTK maintains sensitivity for arbitrarily shaped and scaled waveforms but accounts for the experimental uncertainty when comparing measurements. We demonstrated that the method provides improved accuracy in identifying simulated rhythmic time series and improved consistency across related experimental datasets. More generally, we expect the framework that we have built around JTK_CYCLE [14, 16]—empirical estimation of p-values to account properly for multiple hypothesis testing, analytical approximation of the null distribution, variance “shrinkage” and stabilization, and bootstrapping—can be applied to other rhythm detection algorithms to obtain inexpensive and accurate p-values as we do here.

Our method uses replicates to estimate the variance in expression, which is then propagated to the rhythmicity estimate. For time series data where replicate time points do not exist, a differentapproach is needed. We suggest that the standard deviation of arrhythmic time series is a reasonable approximation of the standard deviation of the time points of rhythmic time series. In Fig. S11, we show that the mean of the standard deviation of the arrhythmic (p> 0.8) time series overestimates the standard deviation for the Hughes *etal.* dataset sampled every 1 h [13] by a factor of about 1.5. While ideally replicate time points would be collected experimentally, we suggest that this approximation be used when replicates are lacking.

**Table 1:** Select functional annotations from the DAVID webtool [11] for the 119 genes identified as rhythmic in 9 or more tissues by applying BooteJTK with a Benjamini-Hochberg adjusted p-value threshold of 0.05 to the Zhang *et al.* dataset [37]. Fold Enrichment refers to how enriched the functional annotation is in the set of genes relative to the expectation for a set of randomly selected genes. Abbreviations: BH, Benjamini-Hochberg adjusted p-value; SP-PIR, Swiss-Prot Protein Information Resource keywords; GO, Gene Ontology keywords.

| Functional Annotation | Fold Enrich. | BH | Genes |
| --- | --- | --- | --- |
| GO:0048511:<br>rhythmic process | 22.57 | 2.11e-11 | ARNTL, CLOCK, CRY1, DBP, HLF, KDR, MMP14, NFIL3, NPAS2, NR1D1, PER1, PER2, PER3, TEF |
| GO:0048770:<br>pigment granule | 16.34 | 2.06e-04 | HSP90AA1, HSPA5, HSPA8, MMP14, PDIA3, PDIA4, SLC1A5 |
| SP-PIR<br>stress response | 24.54 | 3.56e-04 | CIRBP, HSP90AA1, HSPA5, HSPA8, HSPB1, HSPH1 |
| SP-PIR<br>endoplasmic reticulum | 4.07 | 8.68e-04 | CALR, DGAT2, EPHX1, FMO1, FMO2, HERPUD1, HSPA5, LPIN1, P4HA1, PDIA3, PDIA4, POR, SCD2, SDF2L1, SERP1 |
| SP-PIR<br>transcription regulation | 2.62 | 2.52e-03 | ARNTL, BHLHE40, BHLHE41, CLOCK, CRY1, DBP, ELK3, HLF, KLF9, KLF15, LEO1, LITAF, LPIN1, NFIL3, NPAS2, NR1D1, NR1D2, PER1, PER2, PER3, TEF, THRA |
| IPR013126:<br>Heat shock protein 70 | 48.93 | 4.61e-02 | HSPA5, HSPA8, HSPH1 |

Our analysis of the mouse liver microarray dataset from Hughes *et al.* [13] emphasizes the importance of sampling time series frequently and understanding the expected consistency of results. While BooteJTK, in contrast to eJTK, was still able to detect a small fraction of rhythmic genes, even downsampling the data to measurements every 2 h resulted in some differences in genes identified as rhythmic from odd-hour measurements and even-hour measurements. That said, sampling every 2 h is much better than the commonly used 4 h sampling rate, especially when comparisons are made across tissues or conditions. Comparing the tissue overlap results to the 2 h-downsampled benchmark suggests that the aorta tissue samples of Zhang *et al.* [37] were potentially contaminated by brown fat. Future studies can also use the values in Fig. 3 and their analogs for other rhythm detection methods to better understand the overlap and consistency that should be expected when comparing datasets.

Multiple studies have discussed the tissue-specificity of circadian rhythms [23, 29, 37]. Our analysis with BooteJTK supports these claims but suggests that a slightly greater fraction of genes are rhythmic in multiple tissues than was previously appreciated (positive bars at tissue numbers 8 to 12 in Fig. 5B). One limitation of current rhythm detection methods (including eJTK and BooteJTK) is that the null hypothesis states that we have no prior belief or knowledge about the rhythmicity of any given gene. However, genes such as Arntl (Bmal), Nr1d1 (Rev-erbA), Nr1d2 (Rev-erbB), Dbp, Per1, Per2, and Per3 are well-established as circadian and are identified here as rhythmic in all 12 tissues. Given that data for multiple tissues are now available, we can begin to think about how they can be integrated. Simply combining BooteJTK p-values via Fisher’s integration method, which would ignore all tissue-specific knowledge to be gained from sampling 12 tissues, results in 14,598 probes with a Benjamini-Hochberg p-value below 0.05. This provides a pooled estimate for the total number of rhythmic probes in the mouse, ignoring tissue-specific effects.

Methods have been recently developed, however, that can combine information across tissues while allowing for heterogeneity in the context of identifying single-nucleotide polymorphisms (SNPs) that are expression quantitative trait loci (eQTLs) [9]. These methods tend to increase the number of eQTLs that are common to multiple tissues, relative to single-tissue approaches. We expect that applying a similar strategy in rhythm detection would have an analogous effect. For example, Npas2, Hlf, and Tef are identified as rhythmic in 11 of the 12 tissues profiled. Given that these are documented circadian genes [33], it is more likely that these genes are rhythmic in the tissue in which they have not been identified as rhythmic than their p-values suggest. Therefore the current estimates of genes that are rhythmic across several tissues may be underestimates. By the same token, one might doubt that a gene that is rhythmic in only one tissue is genuinely cycling, given the evidence from the other 11 tissues. We were able to address this concern in part by corroborating our results with ChIP-Seq data for core clock transcription factors [20], with the caveat that the ChIP-Seq data was only performed in the mouse liver. Given the increasing amounts of data now available, future methods should systematically integrate multiple sources and types of evidence for rhythm detection.

## 5 Acknowledgements

We would like to thank Matthew Stephens for many discussions regarding bootstrapping and John Hogenesch for sharing data from Zhang *et al.* [37]. This work was completed in part with resources provided by the University of Chicago Research Computing Center.

### Funding

This work was supported by the Defense Advanced Research Projects Agency(D12AP00023) www.darpa.mil/. ALH is a trainee of the National Institutes of Health Medical Scientist Training program at the University of Chicago (grant NIGMS T32GM07281) www.nigms.nih.gov/and is supported in part by the National Institute of Biomedical Imaging And Bioengineering of the National Institutes of Health (Award Number T32EB009412) www.nibib.nih.gov/. The content is solely the responsibility of the authors and does not necessarily reflect the position or the policy of the Government, and no official endorsement should be inferred. The funders had no role in study design, data collection and analysis, decision to publish, or preparation of the manuscript.

## 6 Supplementary Figures

**Figure S1:**
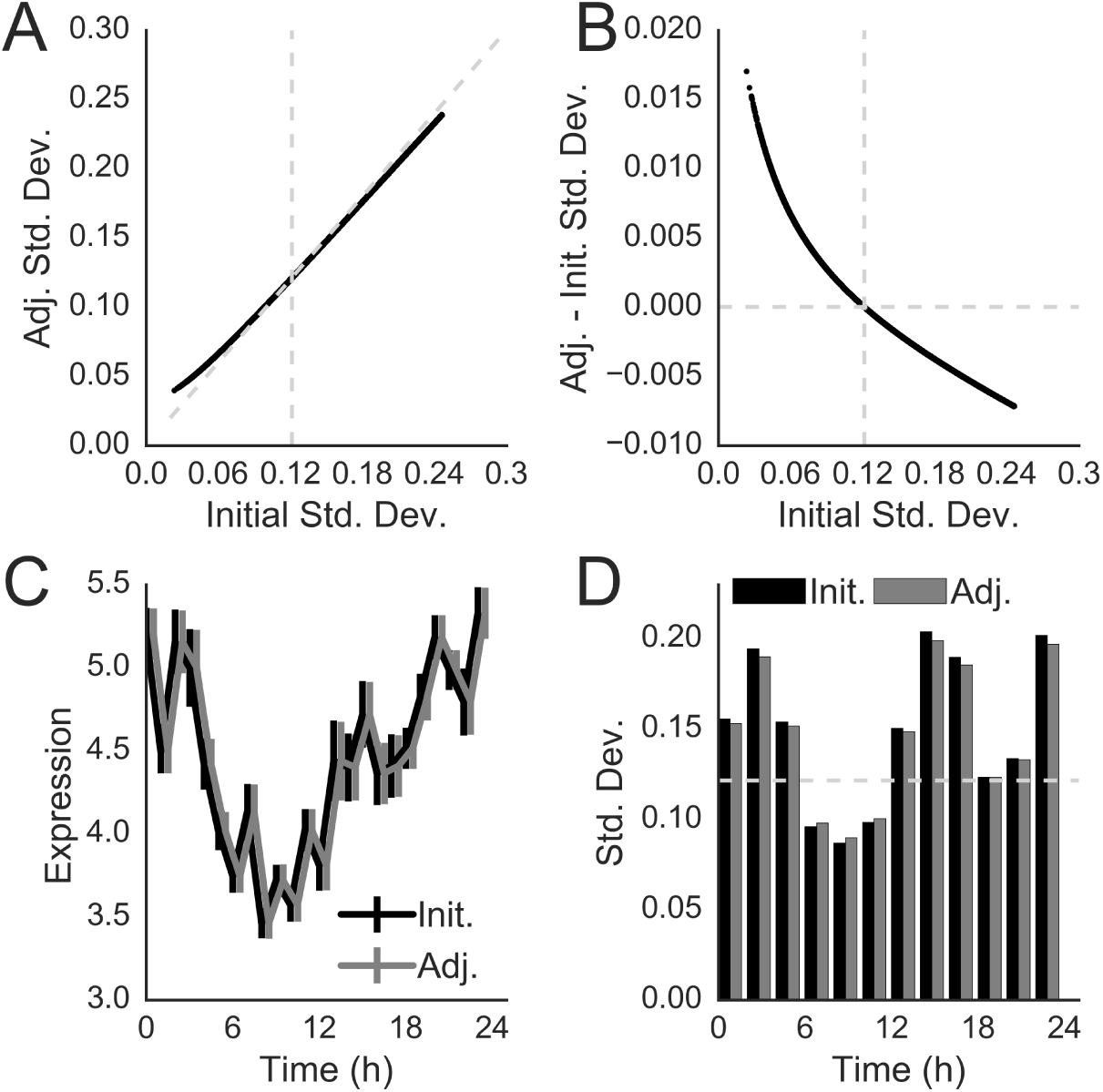
Illustration of the empirical Bayes method. (A) Empirical Bayes adjustment of RNA expression from Hughes *etal.* [13]. Empirical Bayes adjusts the standard deviation estimates to ‘shrink’ the distribution towards the global estimate of 0.1214 (vertical line). The diagonal line has a slope of 1; points below it have been reduced by the eBayes method while points above it have been increased. Shrinkage was performed using *voom* [26] and *vash* [21] in R. (B) The difference between adjusted standard deviation and initial standard deviation is shown against the initial standard deviation. (C) Time series of Clock expression from Hughes *etal.* shown with initial and adjusted standard deviations as error bars. (D) Initial and adjusted standard deviations for Clock RNA expression for even time points from (C).

**Figure S2:**
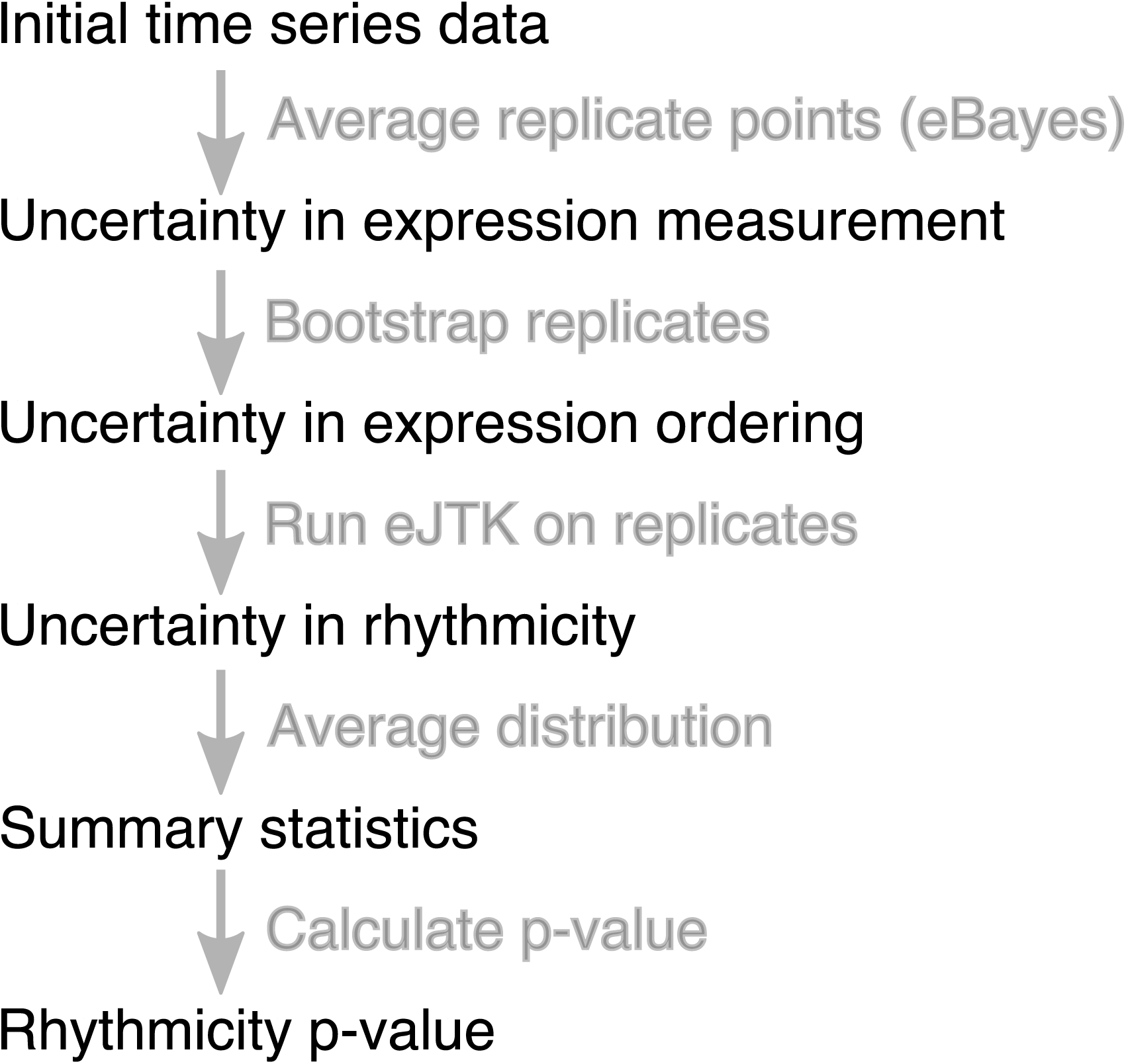
Flowchart of steps in BooteJTK. First, the replicate time points of the initial time series are averaged and standard deviations are obtained via an empirical Bayes (eBayes) procedure. eJTK is then run on bootstrap replicates of the time series. The distribution of eJTK rhythmicity scores are averaged to produce a summary statistic, from which a p-value is calculated.

**Figure S3:**
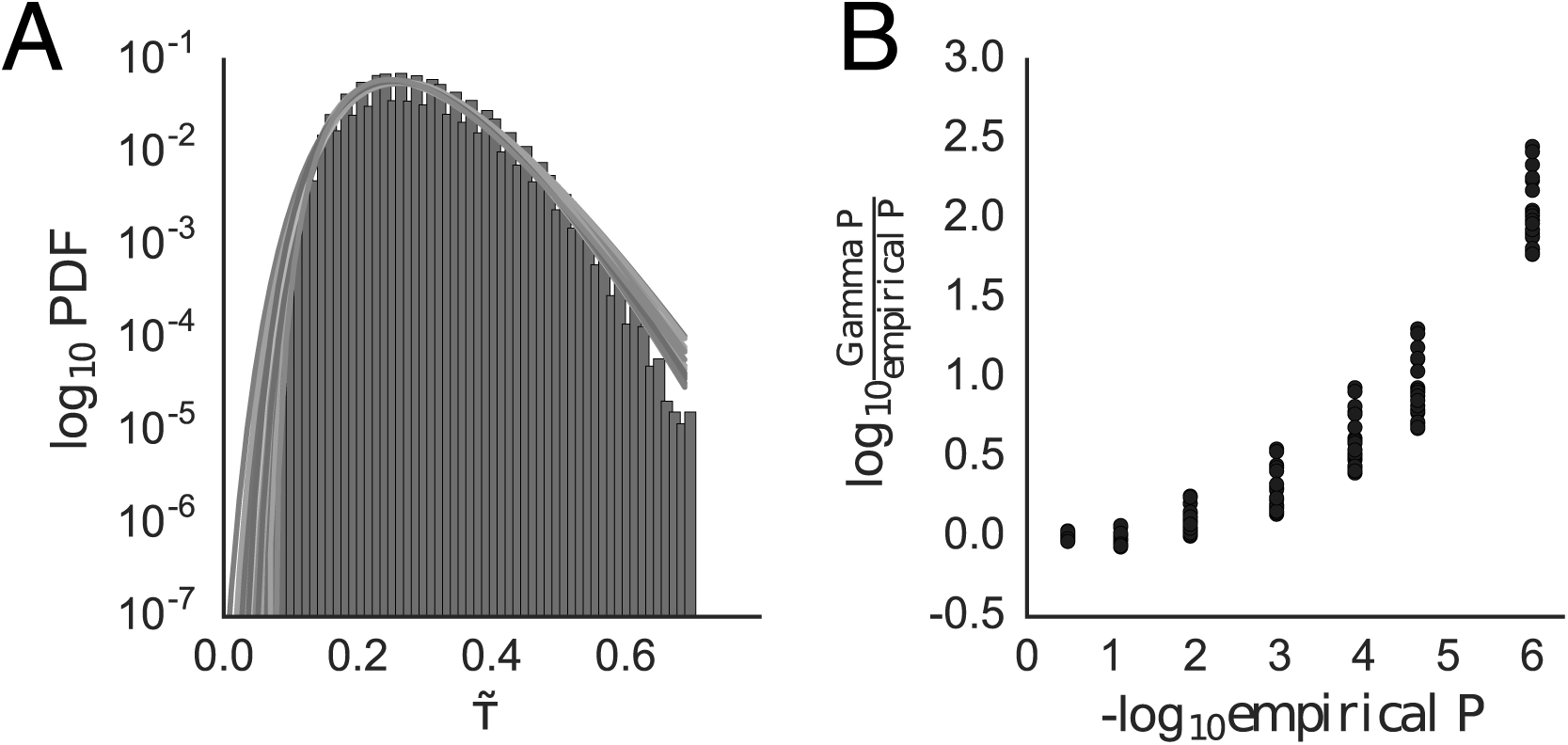
The eJTKτnull distribution can be modeled by a Gamma distribution. (A) Comparison between Gamma distributions parameterized by the Python scipy.stats.gamma.fit [17] function of τ&#x02DC;values for 10^3^ time series comprised of 24 time points drawn randomly from a Gaussian distribution and a histogram of 10^6^τ &#x02DC; values from similarly generated time series. 10 different such Gamma distributions are shown. (B) Logarithms of the ratios of the p-values estimated from the Gamma distributions in (A) (Gamma P) to the p-value calculated directly from the cumulative distribution function for the 10^6^τ &#x02DC; values used to construct the histogram (empirical P). This figure in similar to Fig. S3, which is for BooteJTK.

**Figure S4:**
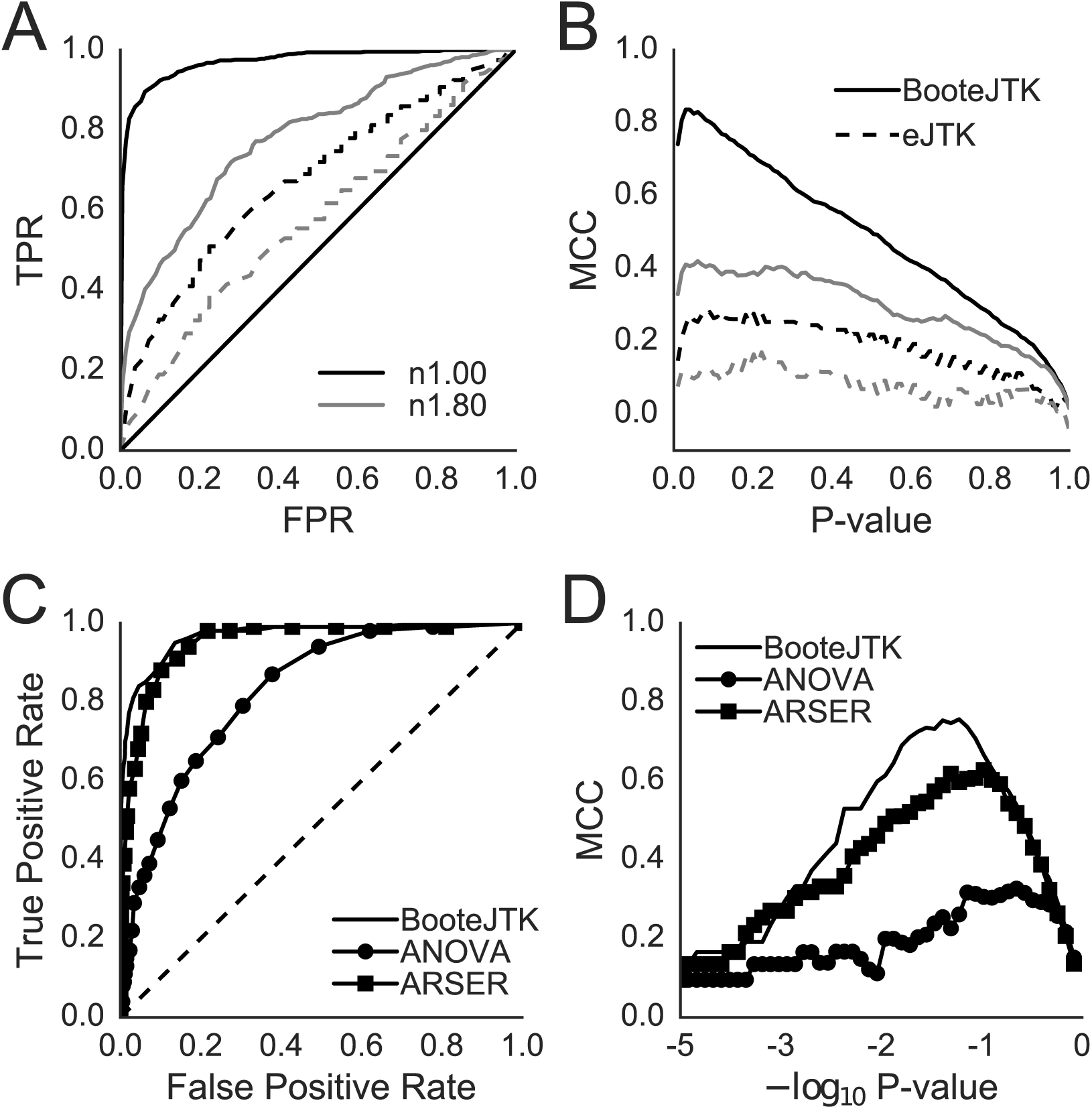
BooteJTK outperforms eJTK, ANOVA, and ARSER. (A & B) Analysis of the effect of noise on algorithmic performance for BooteJTK (solid lines) and eJTK (dashed lines); the noise-to-amplitude ratios are 1.00 (black) and 1.80 (gray). (A) True Positive Rate (TPR) vs. False Positive Rate (FPR) and (B) Matthews Correlation Coefficient (MCC). (C & D) Comparison of algorithmic performance for detecting non-sinusoidal waveforms. Non-sinusoidal series are generated by distorting a cosine so that the time from peak to trough is 4 h and the time from trough to peak is 20 h and adding Gaussian noise with standard deviation that is equal to the amplitude. Algorithms considered are BooteJTK (no symbols), ANOVA (circles), and ARSER (squares) for (C) TPR vs. FPR and (D) MCC.

**Figure S5:**
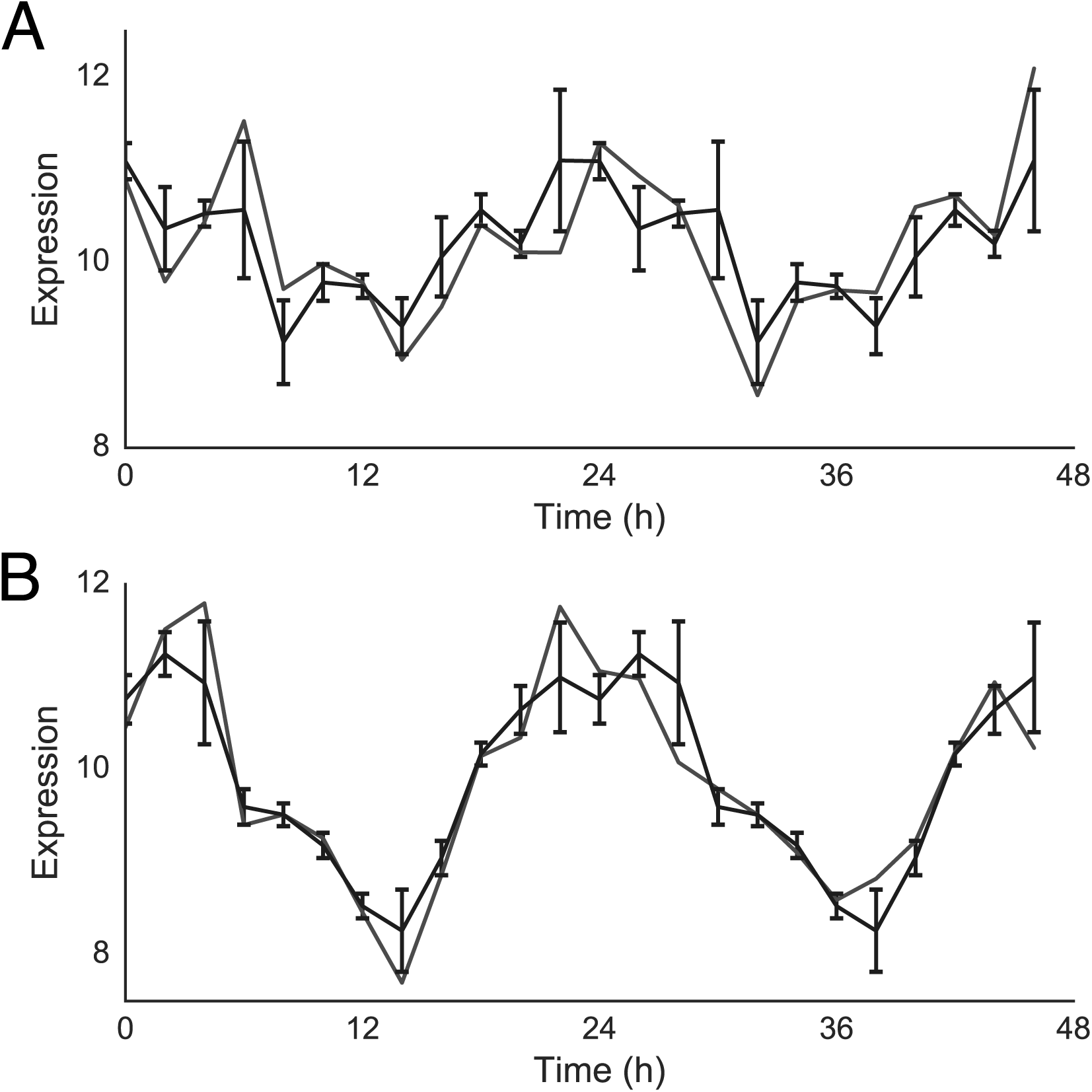
(A) and (B) Examples of two time series with the same eJTKτ &#x02DC; but different BooteJTKτ &#x02DC; values. The grey line is the original time series, the black is the averaged time series with error bars, double plotted to the length of the original time series. The BooteJTKτ &#x02DC; values for (A) and (B) are 0.66 and 0.97 which correspond to p-values of 10^-3^ and 10^-6^. The eJTKτ &#x02DC; score was τ &#x02DC;= 0.57, which corresponds to a p-value of 0.002.

**Figure S6:**
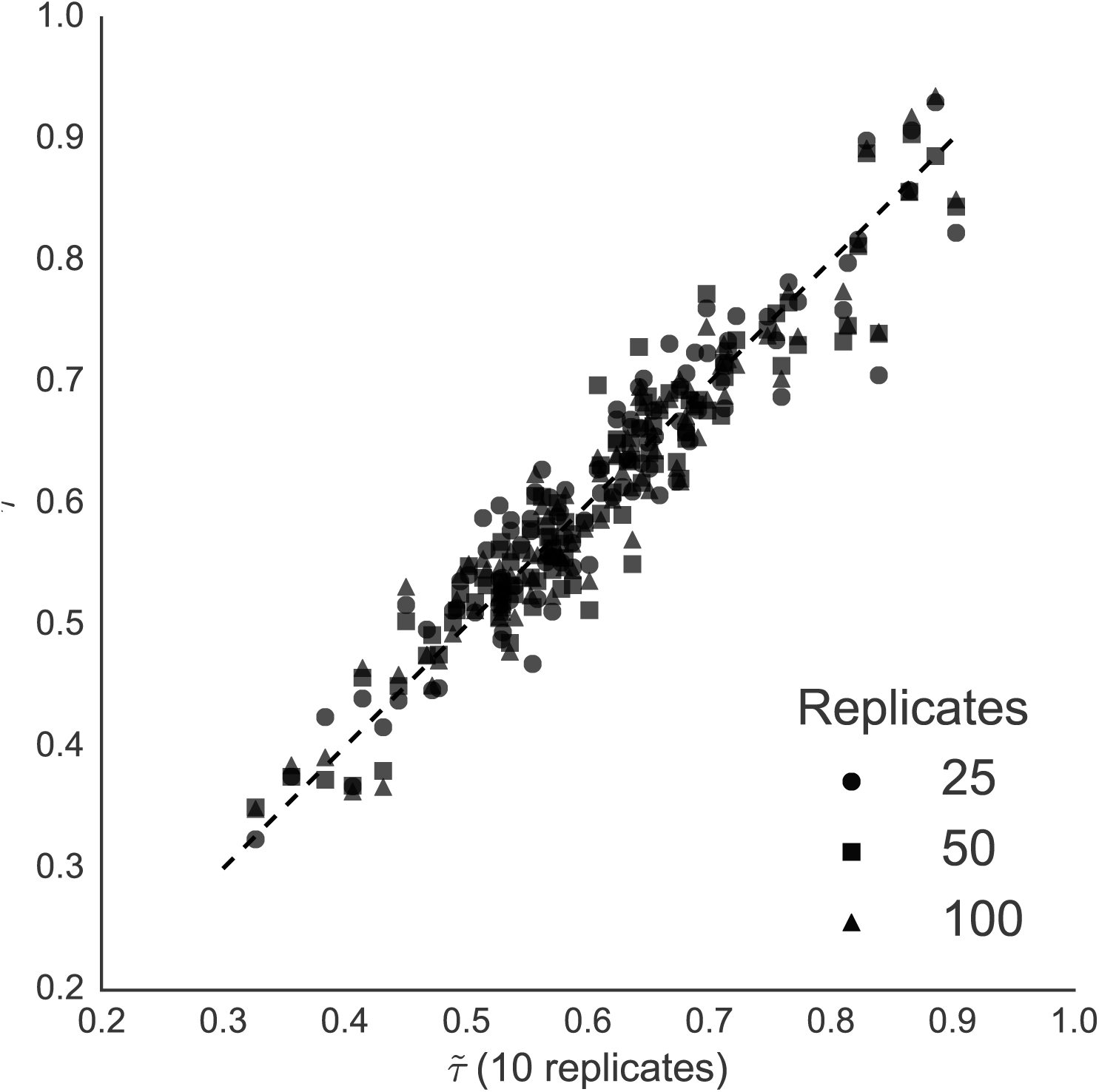
The BooteJTKτ &#x02DC;scores for the same time series dataset show no substantial difference for 10, 25, 50, or 100 bootstrap samples. The diagonal line has a slope of 1.

**Figure S7:**
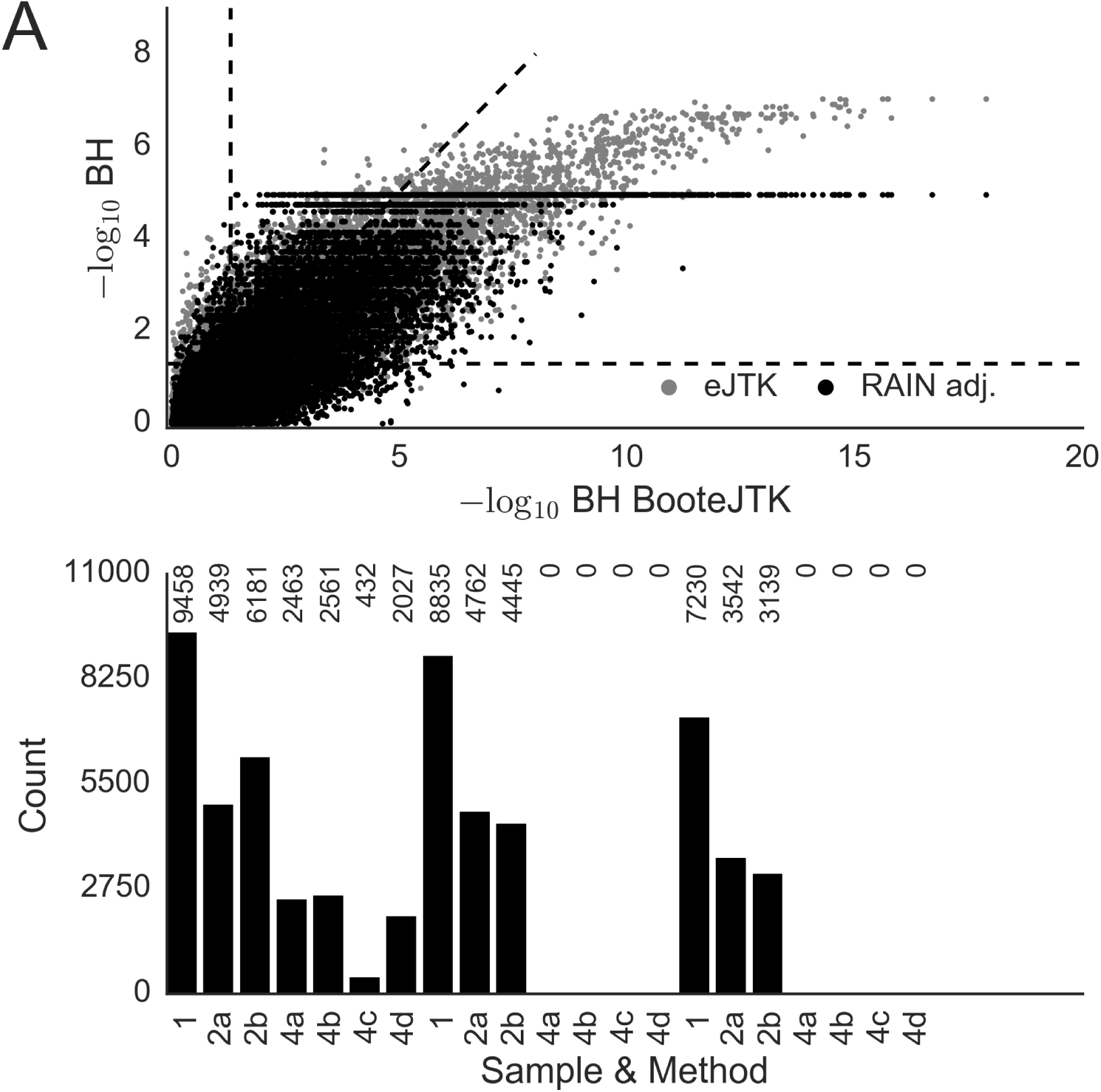
BooteJTK identifies rhythmic probes more consistently than eJTK and RAIN as data are downsampled. Data shown are from Hughes *et al.* 2009 [13] and are originally sampled every 1 h for 48 h; the datasets are downsampled to measurement intervals of 2 h (denoted 2a and 2b) and 4h (denoted 4a, 4b, 4c, and 4d). (A) Comparison of BooteJTK (B) and eJTK (eJ) BH values. Diagonal dashed line has a slope of 1; the horizontal and vertical dashed lines indicate BH = 0.05, as–log_10_(0.05) ≈ 1.3. (B) Number of rhythmic probes at BH < 0.05 for different methods and downsamplings.

**Figure S8:**
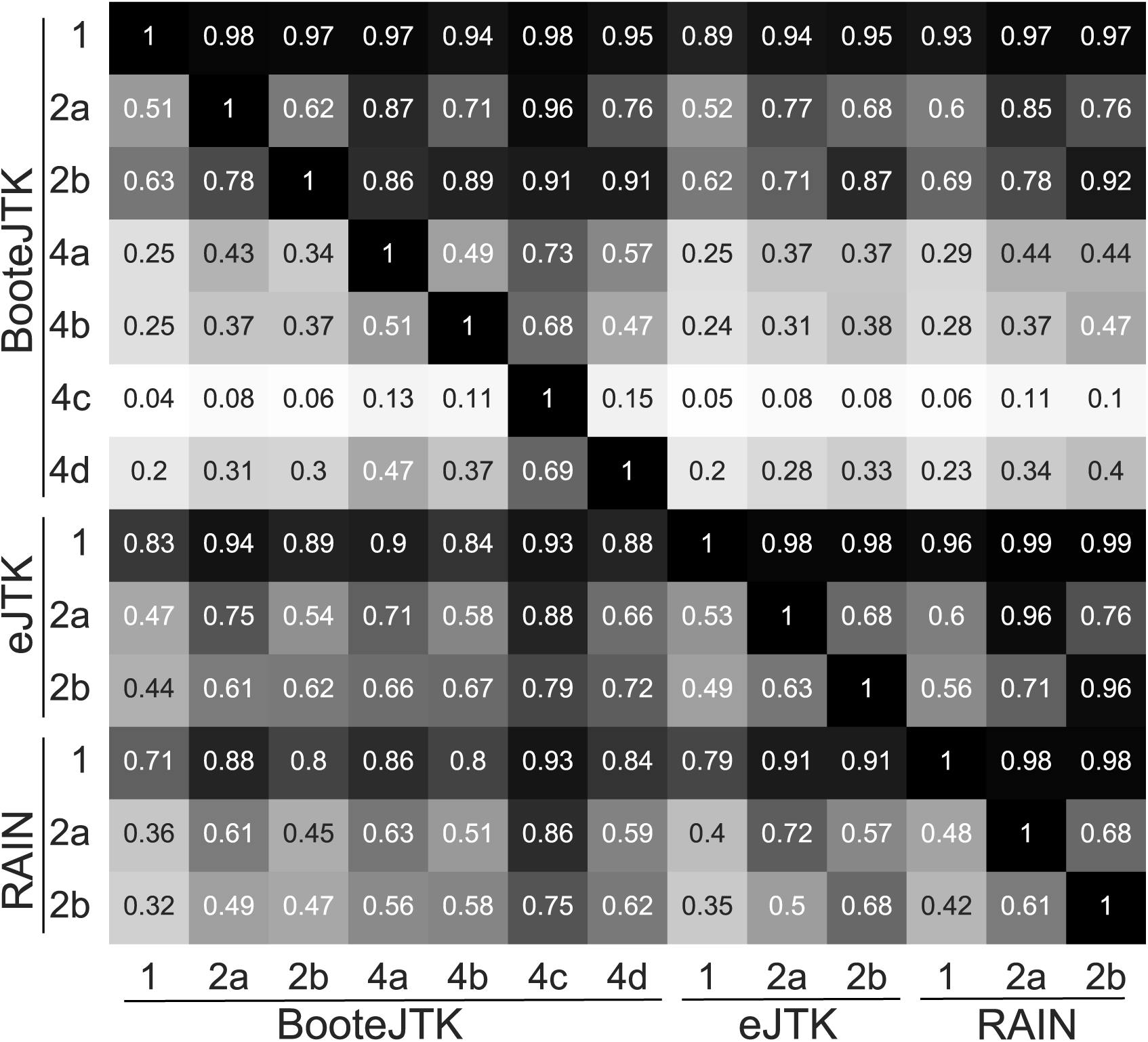
BooteJTKprovides more consistent rhythm detection than eJTK between downsampled datasets. We quantified the overlap between results with different levels of downsampling by the probability that a probe is rhythmic in one dataset (a row) if it is rhythmic in another (a column). As no probes are found to be rhythmic when using eJTK or RAIN on data downsampled to every 4 h, rows and columns for those datasets are not shown. Figure is same as Fig. 4 with RAIN results added.

**Figure S9:**
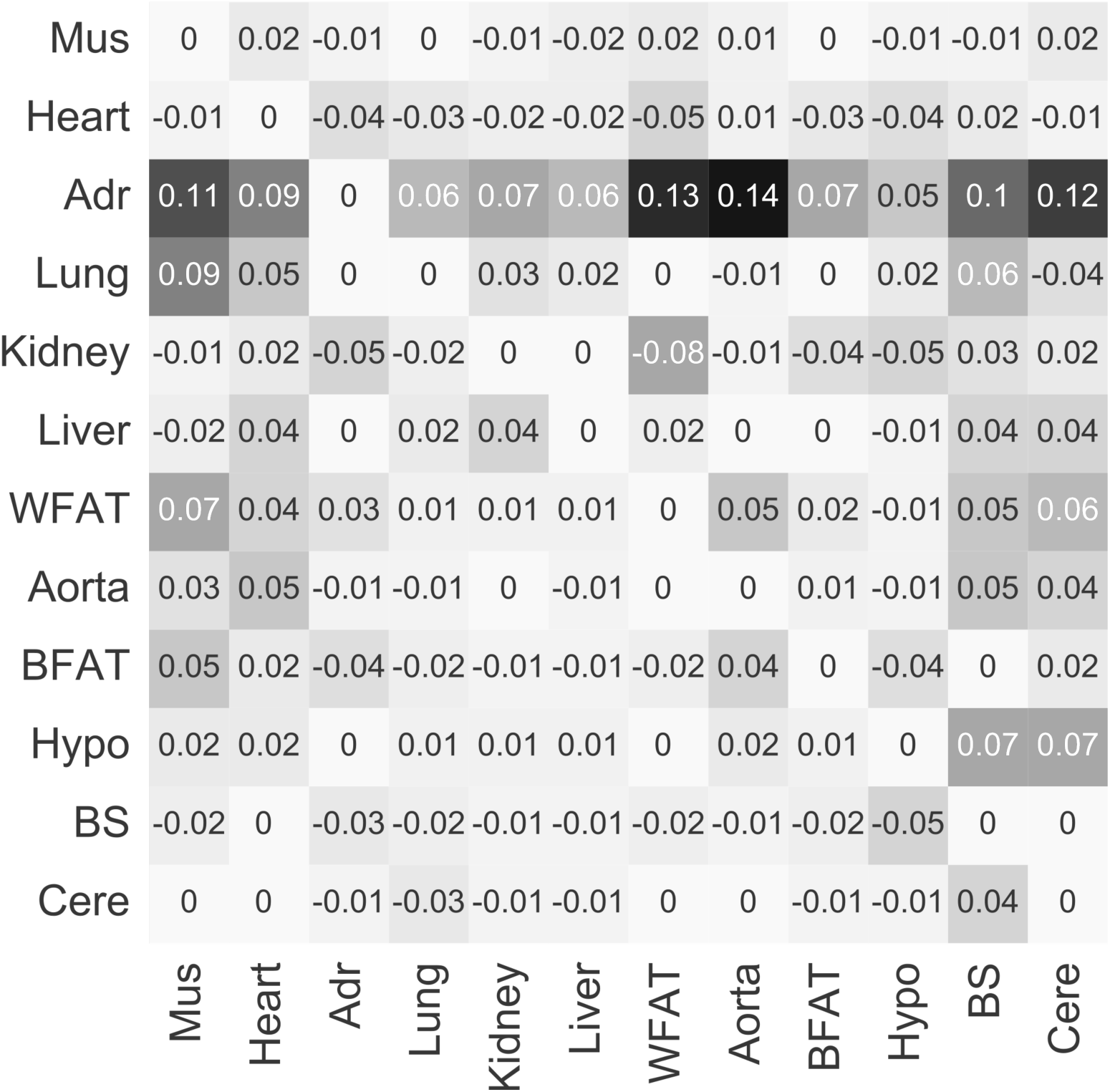
The difference in π probabilities from BooteJTK to eJTK. A positive value indicates a higher π for BooteJTK than eJTK. Tissues are clustered by column π vectors from BooteJTK.

**Figure S10:**
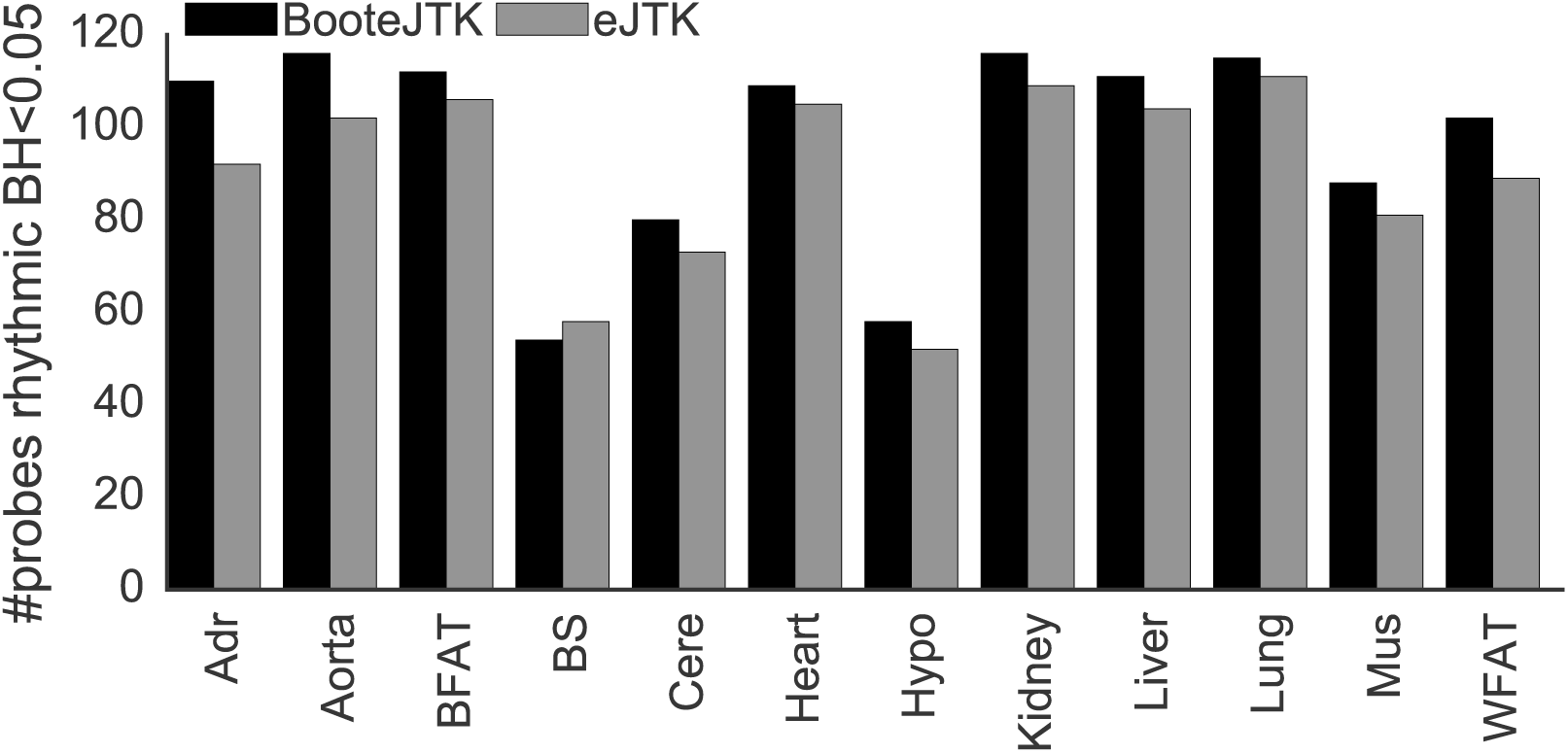
Number of probes rhythmic in each tissue out of the 78 probes rhythmic in 9 or more
tissues. Of the probes rhythmic in more than 9 tissues, many of them are not found to be rhythmic in the brain tissues.

**Figure S11:**
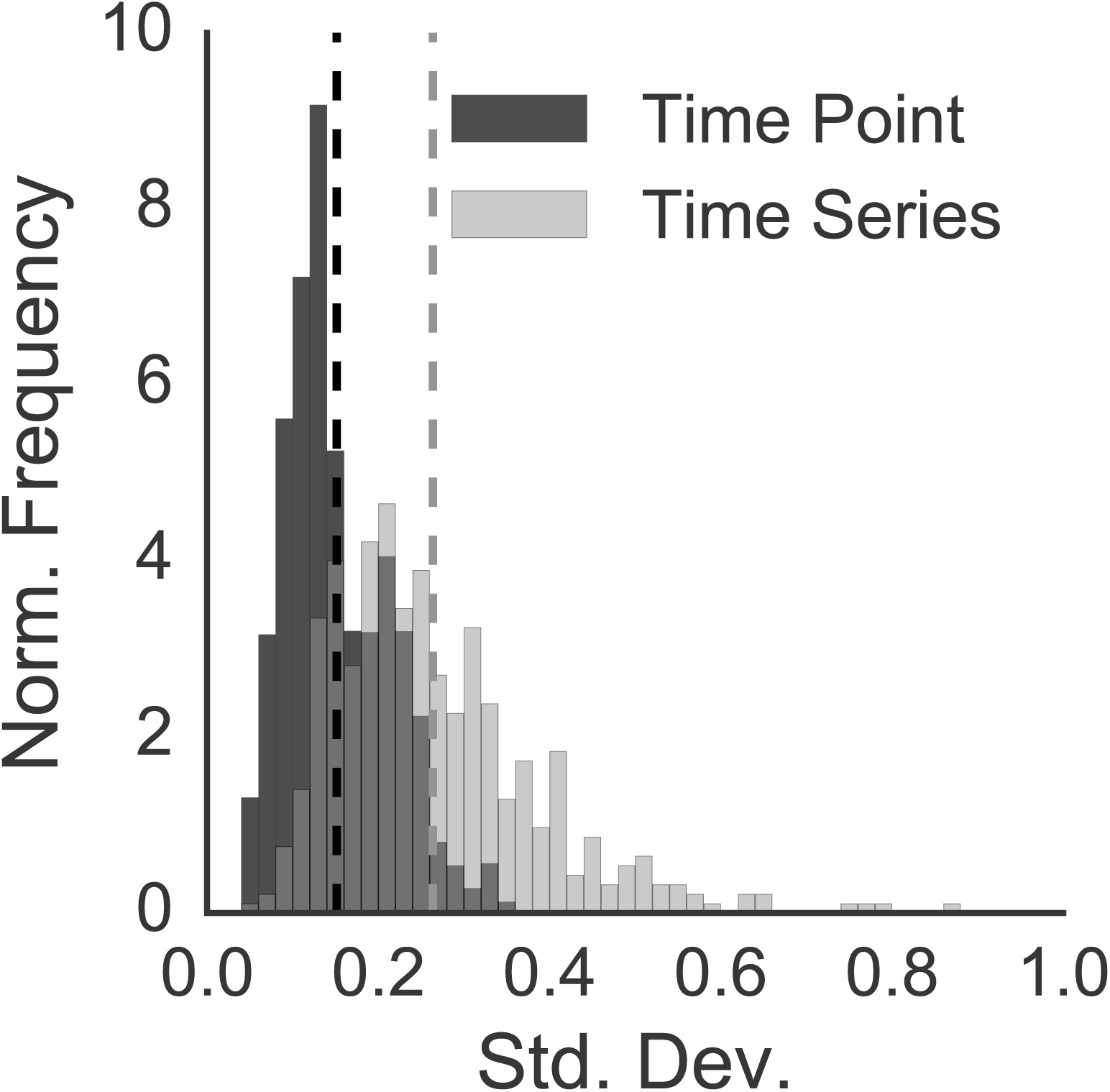
The standard deviation of arrhythmic time series provides an approximation of the
standard deviation of time points, as shown for the Hughes *et al*. dataset sampled every 1 h [13].
Vertical lines represent the means of the two distributions.

## References

[1]. Yoav Benjamini and Yosef Hochberg. Controlling the False Discovery Rate: A Practical and Powerful Approach to Multiple Testing. Journal of the Royal Statistical Society, Series B (Methodological), 571:289–300, (1995).

[1] Nicolas L Bray, Harold Pimentel, Pall Melsted, Lior Pachter. Near-optimal probabilistic RNA-seq quantification. Nat Biotech, 345:525–527, May (2016).

[3] Morton B Brown. A Method for Combining Non-Independent, One-Sided Tests of Significance. Biometrics, 314:987–992, (1975).

[4] Anastasia Deckard, Ron C Anafi, John B Hogenesch, Steven B Haase, and John Harer. Design and analysis of large-scale biological rhythm studies: a comparison of algorithms for detecting periodic signals in biological data. Bioinformatics, 2924:3174–80, Dec (2013).

[5] Bradley Efron and Robert J Tibshirani. An Introduction to the Bootstrap. CRC press, (1994).

[6] R A Fisher. Statistical Methods for Research Workers. Oliver and Boyd, (1925).

[7] Timothy P Fitzgibbons, Sophia Kogan, Myriam Aouadi, Greg M Hendricks, Juerg Straubhaar, and Michael P Czech. Similarity of mouse perivascular and brown adipose tissues and their resistance to diet-induced inflammation. American, Journal of Physiology. Heart and Circulatory Physiology, 3014:H1425–37, Oct 2011.

[8] Matthieu Flourakis, Elzbieta Kula-Eversole, Alan L. Hutchison, Tae Hee Han, Kimberly Aranda, Devon L. Moose, Kevin P. White, Aaron R. Dinner, Bridget C. Lear, Dejian Ren, Casey O. Diekman, Indira M. Raman, and Ravi Allada. A Conserved Bicycle Model for Circadian Clock Control of Membrane Excitability. Cell, 1624:836–848, (2015).

[9] Timothée Flutre, Xiaoquan Wen, Jonathan Pritchard, and Matthew Stephens. A Statistical Framework for Joint eQTL Analysis in Multiple Tissues. PLoS Genetics, 95:1–8, (2013).

[10] Akihiro Goriki, Fumiyuki Hatanaka, Jihwan Myung, Jae Kyoung Kim, Takashi Yoritaka, Akio Matsubara, Daniel Forger, and Toru Takumi. A Novel Protein, CHRONO, Functions as a Core Component of the Mammalian Circadian Clock. PLoS Biology, 124: (2014).

[11] Da Wei Huang, Brad T Sherman, and Richard A Lempicki. Bioinformatics enrichment tools: paths toward the comprehensive functional analysis of large gene lists. Nucleic Acids Research, 371:1–13, Jan (2009).

[12] Da Wei Huang, Brad T Sherman, and Richard A Lempicki. Systematic and integrative analysis of large gene lists using DAVID bioinformatics resources. Nature Protocols, 41:44–57, Jan (2009).

[13] Michael E. Hughes, Luciano DiTacchio, Kevin R. Hayes, Christopher Vollmers, S. Pulivarthy, Julie E. Baggs, Satchidananda Panda, and John B. Hogenesch. Harmonics of Circadian Gene Transcription in Mammals. PLoS Genet, 54:e1000442, Apr (2009).

[14] Michael E Hughes, John B Hogenesch, and Karl Kornacker. JTK_CYCLE: an efficient nonparametric algorithm for detecting rhythmic components in genome-scale data sets. Journal of Biological Rhythms, 255:372–80, Oct (2010).

[15] Alan L. Hutchison and Aaron R. Dinner. Correcting for Dependent P-values Improves Accuracy of Leading Rhythm Detection Methods. Inpreparation (2017).

[16] Alan L. Hutchison, Mark Maienschein-Cline, Andrew H. Chiang, S. M Ali Tabei, Herman Gudjonson, Neil Bahroos, Ravi Allada, and Aaron R. Dinner. Improved Statistical Methods Enable Greater Sensitivity in Rhythm Detection for Genome-Wide Data. PLoSComputBiol, 113:e1004094, (2015).

[17] Eric Jones, Travis Oliphant, and Pearu Peterson. Scipy: Open source scietific tools for Python, (2001).

[18] Kevin P. MKeegan, Suraj Pradhan, Ji Ping Wang, and Ravi Allada. Meta-analysis of Drosophila circadian microarray studies identifies a novel set of rhythmically expressed genes. PLoS Computational Biology, 311:2087–2110, (2007).

[19] Douglas R Kellogg. Wee1-dependent mechanisms required for coordination of cell growth and cell division. Journal of Cell Science, 11624:4883–4890, Nov (2003).

[20] Nobuya Koike, Tae-kyung Kim, and Joseph S Takahashi. Transcriptional Architecture and Chromatin Landscape of the Core Circadian Clock in Mammals. Science, 338(August):1–10, (2012).

[21] Mengyin Lu and Matthew Stephens. Variance Adaptive Shrinkage (vash): Flexible Empirical Bayes estimation of variances. bioRxiv, (July):048660, (2016).

[22] Takuya Matsuo, Shun Yamaguchi, and Shigeru Mitsui. Control Mechanism of the Circadian Clock for Timing of Cell Division In Vivo. Science, 302(October):255–260, (2003).

[23] Satchidananda Panda, Marina P. Antoch, Brooke H. Miller, Andrew I. Su, Andrew B. Schook, Marty Straume, Peter G. Schultz,Steve A. Kay, Joseph S. Takahashi, and John B. Hogenesch. Coordinated Transcription of Key Pathways in the Mouse by the Circadian Clock. Cell, 1093:307–320, (2002).

[24] Mark Perelis, Biliana Marcheva, Kathryn Moynihan Ramsey, Matthew J. Schipma, Alan L. Hutchison, Akihiko Taguchi, Clara Bien Peek, Heekyung Hong, Wenyu Huang, Chiaki Omura, Amanda L. Allred, Christopher A. Bradfield, Aaron R. Dinner, Grant D. Barish, and Joseph Bass. Pancreatic *β* cell enhancers regulate rhythmic transcription of genes controlling insulin secretion. Science, 3506261:aac4250, (2015).

[25] Harold J Pimentel, Nicolas Bray, Suzette Puente, Páll Melsted, and Lior Pachter. Differential analysis of RNA-Seq incorporating quantification uncertainty. bioRxiv, Jun (2016).

[26] Matthew E Ritchie, Belinda Phipson, Di Wu, Yifang Hu, Charity W Law, Wei Shi, and Gordon K Smyth. limma powers differential expression analyses for RNA-sequencing and microarray studies. Nucleic Acids Research, 437:e47, (2015).

[27] Oded Sandler, Sivan Pearl Mizrahi, Noga Weiss, Oded Agam, Itamar Simon, and Nathalie Q Balaban. Lineage correlations of single cell division time as a probe of cell-cycle dynamics. Nature,5197544:468–471, Mar (2015).

[28] Gordon K Smyth. Linear Models and Empirical Bayes Methods for Assessing Differential Expression in Microarray Experiments Linear Models and Empirical Bayes Methods for Assessing Differential Expression in Microarray Experiments. Statistical Applications in Genetics and Molecular Biology Volume, 31:1–26, (2004).

[29] Kai-Florian Storch, Ovidiu Lipan, Igor Leykin, N Viswanathan, Fred C Davis, Wing H Wong, and Charles J L B Weitz. Extensive and divergent circadian gene expression in liver and heart. Nature, 4176884:78–83, (2002).

[30] Paul F Thaben and PålO Westermark.Detecting rhythms in time series with RAIN. Journal of Biological Rhythms, 296:391–400, (2014).

[31] Cole Trapnell, Adam Roberts, Loyal Goff, Geo Pertea, Daehwan Kim, David R Kelley, Harold Pimentel, Steven L Salzberg, John L Rinn, and Lior Pachter. Differential gene and transcript expression analysis of RNA-seq experiments with TopHat and Cufflinks. Nature Protocols, 73:562–78, Mar (2012).

[32] John W Tukey. Exploratory Data Analysis. Addison-Wesley, (1977).

[33] Hideki Ukai and Hiroki R Ueda. Systems Biology of Mammalian Circadian Clocks. Annual Peview of Physiology., 72:579–603, (2010).

[34] Gang Wu, Ron C Anafi, Michael E Hughes, Karl Kornacker, and John B Hogenesch. Meta-Cycle: an integrated R package to evaluate periodicity in large scale data. Bioinformatics, 1-3(July):040345, (2016).

[35] Jean Wu, James MacDonald,Jeff Gentry, and Rafael Irizarry. gcrma: Background Adjustment Using Sequence Information, (2016).

[36] Rendong Yang and Zhen Su. Analyzing circadian expression data by harmonic regression based on autoregressive spectral estimation. Bioinformatics, 2612: (2010).

[37] Ray Zhang, Nicholas F Lahens, Heather I Ballance, Michael E Hughes, and John B Hogenesch. A circadian gene expression atlas in mammals: implications for biology and medicine. Proceedings of the National Academy of Sciences, 11145:16219–24, (2014).

